# Arm Control and its Recovery after Selective Lesions of Sensorimotor Cortex and the Red Nucleus: A Kinematic Study in Non-Human Primates

**DOI:** 10.1101/2025.08.06.668715

**Authors:** Anna Baines, Annie Poll, Anne M.E. Baker, John W. Krakauer, Stuart N. Baker

**Affiliations:** Faculty of Medical Sciences, Newcastle University, Newcastle upon Tyne, NE2 4HH, UK; Departments of Neurology and Neuroscience, Johns Hopkins University, Baltimore, MD, USA; The Santa Fe Institute, Santa Fe NM, USA

## Abstract

Cortical and subcortical lesions to the motor system, as often occur with stroke, typically lead to transitions through a stereotyped upper limb recovery sequence. After initial weakness and loss of dexterity, spasticity and fixed muscle coactivation patterns (synergies) appear. Early work suggested that different features arise from distinct primary motor cortex (M1) subdivisions. Here we investigated this with modern methods, using ischemic lesions of various cortical areas and electrocoagulation lesions of magnocellular red nucleus (RNm) in rhesus monkeys.

Nine animals were trained on a reach and grasp task; hand kinematics were assessed with markerless tracking. The proportion of damaged cortical layer V cells in each cortical area was quantified, and corresponding kinematic effects evaluated. Reaching speed showed greater and more persistent reductions with larger lesions to the posterior part of M1 on the gyrus (Posterior Old M1 in Strick’s terminology). Initial increases in trajectory variability were more consistent with greater damage within the central sulcus (New M1); these partially recovered. Lesions involving Anterior Old M1 (Area 4s in Hines’ terminology) had no additive negative effects. An extensive cortical lesion, which combined New and Old M1 with pre-motor and somatosensory cortex damage did not produce a worse or more persistent deficit than lesions limited to M1, suggesting that loss of arm control arose mainly from damage to descending pathways rather than cortico-cortical interactions.

Lesions of RNm led to long-lasting slowing of reach, but no increase in variability. Subsequent cortical lesions to Old M1 led to more severe effects, and worse recovery, than without the preceding RNm lesion. This suggests an important neural compensatory role for the rubrospinal tract following cortical damage in monkey, which is not available in humans where the rubrospinal tract is vestigial.

None of the lesions investigated led to overt abnormal synergies.

The results are consistent with known differences in descending connections from each area: New M1 has fast cortico-motoneuronal output, known to be important for fine motor control (here assessed by trajectory variability); Old M1 has cortico-reticular connections able to activate the reticulospinal tract, important for generating the high forces needed for fast movements.

## Introduction

Damage to the primary motor cortex (M1) and its descending corticofugal projections, as commonly seen after stroke, results in contralateral hemiparesis. This is seen in more than 80% of patients acutely and 40% chronically (Cramer et al., 1997), as most strokes occur in middle cerebral artery (MCA) territory (Shelton and Reding, 2001). Twitchell (1951) and Brunnstrom (1970) described that upper limb recovery after such strokes typically ‘occurs in an almost standardized fashion’ (Brunnstrom, 1966). Initial weakness and loss of dexterity (negative signs) are replaced with involuntary increases in muscle activity (positive signs). Heightened reflex responses emerge into poorly coordinated movements dominated by highly stereotyped muscle coactivation patterns or ‘synergies’. Abnormal synergies usually eventually disappear, making isolated joint movements possible again. However, recovery can stall at any one of these stages (Twitchell, 1951; Brunnstrom, 1970), leaving moderate to severe patients with movement repertoires confined to synergies (Dewald et al., 1995; Dewald and Beer, 2001; Beer et al., 2007; Ellis et al., 2007). In patients with the abnormal flexor synergy, shoulder abduction drives simultaneous activation of elbow, wrist and finger flexors, limiting elbow extension (Sukal et al., 2007; Miller and Dewald, 2012; Lan et al., 2017; Avni et al., 2024). MCA strokes are variable, with infarcts often damaging multiple cortical areas in tandem. This prevents assignment of different features of the hemiparetic phenotype (i.e. weakness, loss of dexterity, spasticity, synergies) to the degree of damage in specific brain regions. Targeted surgical lesions in an animal model paired with careful observation of behavioral effects could provide fundamental knowledge linking anatomy to component deficits. This could allow rational stroke therapies, targeting different treatments based on the degree of damage in each area.

Studies performed in monkey over 80 years ago showed that positive motor signs did not result from pyramidal tract lesions (Tower, 1940), but were seen after lesions to M1 (Hines, 1937, 1943). Hines noted that positive signs were strongest after damage to a strip approximately 3mm wide located just posterior to Brodmann’s area 6 (premotor cortex). Hines reported that this subregion, which she referred to as ‘Area 4s’, primarily suppresses motor output when stimulated; this was in contrast to mainly excitatory effects from stimulating more posterior M1. Suppressive effects from Area 4s may be mediated by corticobulbar connections (McCulloch et al., 1946) to a region of the reticular formation which suppresses motor output (Magoun and Rhines, 1946; Engberg et al., 1968; Jankowska et al., 1968; Takakusaki et al., 2001; Du Beau et al., 2012). The Area 4s designation has fallen out of current usage, although a modern study which mapped output effects from M1 and pre-motor cortex using microstimulation did show a clear band of mainly suppressive effects in the most anterior portion of M1 in one out of the two monkeys examined (Boudrias et al., 2010).

A different approach to subdividing M1 was taken by Rathelot and Strick (2009), who mapped the distribution of cortico-motoneuronal (CM) cells. They reported that CM cells were mainly found in the bank of the central sulcus, not on the gyrus, and coined the terms ‘New M1’ and ‘Old M1’ to refer to these two regions. Both parts of M1 have a similar density of corticospinal neurons (Dum and Strick, 1991; He et al., 1993); only Old M1 connects to the brainstem (Keizer and Kuypers, 1989). New M1 is found only in Old World monkeys and humans, and is considered essential for performing dexterous upper limb movements (Lemon, 2008; Rathelot and Strick, 2009). Witham et al. (2016) provided evidence that Old M1 also has CM cells (as was already clear from the work of Rathelot and Strick (2009)) but that these have more slowly conducting axons than those in New M1. Area 4s of Hines would occupy approximately the anterior half of Old M1 as described by Rathelot and Strick.

Given this extensive anatomical and physiological evidence, we hypothesized that focal lesions to the different M1 subregions would lead to distinct post-lesion phenotypes. However, post-lesion deficits are a composite not only of the function that has been lost, but also of the nervous system’s ability to recover and restore that function. Following mild stroke, spared parts of M1 underlie functional recovery (Nudo and Milliken, 1996; Nudo et al., 1996; Rouiller et al., 1998). After larger M1 lesions, ipsilesional motor regions such as the dorsal premotor cortex (PMd) and the supplementary area (SMA) can be recruited to replace lost corticospinal output (Liu and Rouiller, 1999; Johansen-Berg et al., 2002; Fridman et al., 2004; McNeal et al., 2010). In even larger strokes where these areas are also damaged, there is an increased reliance on the contralesional motor cortex to restore connections to the affected side via the cortico-reticulospinal pathway (Zaaimi et al., 2012; Zaaimi et al., 2018). This compensation partially restores upper limb reaching, albeit with poorer quality movements (McPherson et al., 2018; Choudhury et al., 2025).

Charles Sherrington once said, *‘…the pyramidal tract is such a human feature… in macaque the recovery is surprising, when contrasted with the degree and persistence of the human condition, which is often permanent, is it not?’* (Walshe, 1961). The marked difference in speed and quality of recovery between macaques and humans after a cortical lesion remains an enigma. One possible explanation could come from differences in the rubrospinal pathway. In humans, the magnocellular red nucleus (RNm) outputs to the spinal cord are notably weak, reflecting an evolutionary shift towards greater cortical control of movement (Nathan and Smith, 1955; Onodera and Hicks, 2010; Herculano-Houzel et al., 2016). By contrast, in monkey the rubrospinal tract is an important motor pathway (Cheney, 1980; Holstege et al., 1988; Ralston et al., 1988; Mewes and Cheney, 1991; Riddle et al., 2009), and is known to be involved in functional recovery (Belhaj-Saïf and Cheney, 2000). It is plausible that without a functional rubrospinal system, monkeys would exhibit similarly limited upper limb recovery as seen in humans; however, this remains unconfirmed.

In this study, we set out to test if lesions to the distinct motor cortical subareas described above would result in different deficits and different recovery time courses. We measured movement speed and trajectory variability, considering them the reaching analogues of weakness and dexterity in hand function (Cortes et al., 2017; Hadjiosif et al., 2021). We also examined whether recovery was worsened by prior unilateral lesions to RNm, or by extending cortical lesion size. We show impairments in both movement variability and speed after lesions; trajectory variability was only affected following lesions that damaged New M1 or the RNm; movement slowing was long-lasting after lesions with greater Posterior Old M1 involvement. Lesions of Anterior Old M1 (equivalent to Area 4s) had minimal effects. RNm lesions slowed movement only a little, but recovery from a cortical lesion after a preceding RNm lesion was markedly impaired. None of the focal lesions which we tested generated overt positive signs.

## Materials and Methods

Experiments were conducted in one male (weight at start of study = 11.1 kg) and eight female rhesus macaques (weights at start of study = 5.5-7.7 kg), referenced in this report as Monkey D, B, Ca, Ch, Cm, Co, Cu, Ze, and Zd respectively. All animal procedures were carried out under an appropriate license issued by the UK Home Office under the Animals (Scientific Procedures) Act 1986 and were approved by the Animal Welfare and Ethical Review Board of Newcastle University.

### Behavioral Task

Animals were trained to perform a reach and grasp task with their right upper limb (Fig. 1A). The neck and left limb were gently restrained throughout. A trial of the task was initiated by the monkey holding a handle for one second. The lid of a baited cup then rapidly retracted, providing access to a food reward contained within. The monkey made a rapid reaching movement to retrieve the reward, after which the trainer pressed a foot switch to signal the successful end of the trial. A timeout tone sounded if the footswitch was not pressed within 5 s of handle release; the trial then had to be repeated from the start. Five or ten repeat trials were performed reaching in the same direction, after which the handle and cup were moved to another of the 12 possible arrangements on the vertices of a diamond (Fig. 1B). This allowed for performance of an assortment of reaches requiring elbow flexion or extension each day.

**Figure 1.**
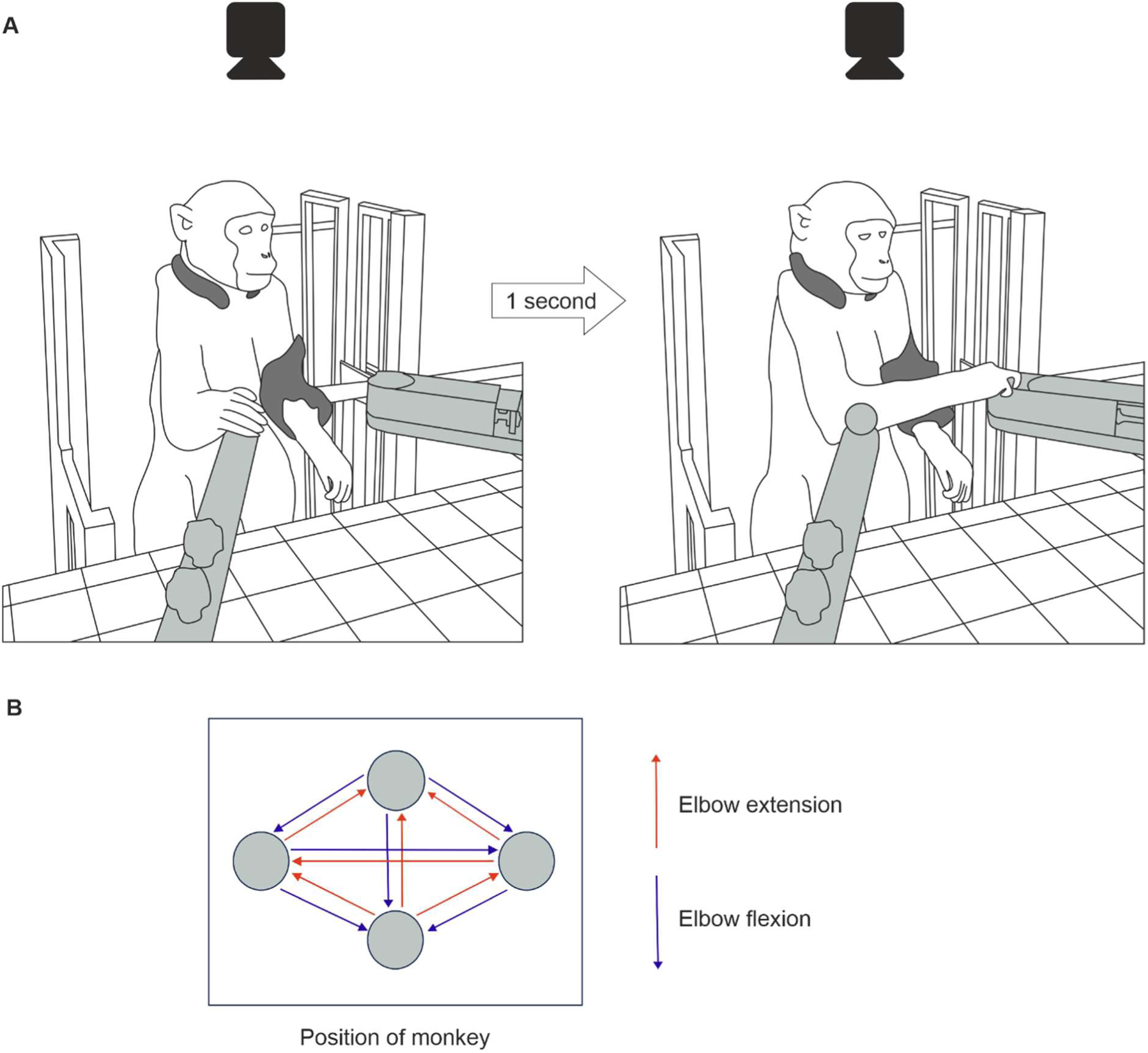
Behavioral task. A, drawings of the reaching task setup. Monkeys were placed in a neck and left arm restraint (dark grey). Handle and cup (light grey) positions were fixed to the table; here the trial has the handle on the animal’s right, and the cup on the left. Holding the handle for one second unlocked the cup, and the food reward was retrieved. A camera positioned above the monkey recorded high speed video footage used for kinematic analysis. B, twelve possible handle and cup arrangements for assessing both elbow extension and flexion reach performance, shown with the same perspective as the camera’s field of view.

Most monkeys learnt this task rapidly (3-6 months), and performed it reliably on each session. Three monkeys (Monkey Ze, B and Cm) were consistently given oral trazodone to improve focus and task performance (25-62.5 mg daily dose given 1 hour before training).

### Video Recording and Markerless Tracking

Baseline data were collected 5 days per week for 2-5 weeks to sample normal reaching behavior, before the lesion was made. Further kinematic data were then gathered for 15 weeks.

High-speed video footage was recorded from a camera positioned above the monkey (90-100 fps; 1280x1024 resolution; CM3-U3-31S4C-CS 1/1.8” Chameleon**®**3 Color Camera, Teledyne FLIR, Oregon, USA). Frame capture by the camera was driven by regular digital trigger pulses, which were generated by a custom circuit containing an Arduino microcontroller. The microcontroller also generated an 8 bit binary counter, which incremented once with each trigger. The counter was connected to 8 LEDs visible within the camera frame. Rarely (typically <1 in 1000 frames), the camera software missed writing a frame to disk. We identified such missed frames using a custom MATLAB script which extracted and thresholded the light intensities around each LED to determine the binary count and detect non-contiguous frames. Videos and task parameters were stored to disk.

Videos were processed offline using the open source markerless tracking software DeepLabCut (DLC, version 2.2, (Nath et al., 2019)). A ResNet-50 model pre-trained on an ImageNet database was custom-trained on task video footage to predict right hand location accurately in each video frame. Initial training frames were extracted using a k-means algorithm, and positions of the handle, cup and right index finger metacarpophalangeal joint were manually labelled in each frame. The network was iteratively trained, and prediction accuracy of the three landmarks was reviewed using test video footage (i.e. frames not used for network training). Any further network training was carried out using additional frames manually chosen from poorer tracking scenarios. 3640 frames from sessions with different animals were used for training in total. The error of different iterations of the trained network used for task video analysis was 2-2.6 mm, testing error 2.4-9.5 mm (measured with 0.6 prediction likelihood cutoff).

Hand position and speed over time were low pass-filtered at 8 Hz and plotted using custom-written MATLAB scripts (Fig. 2). A trial was included in subsequent analyses if it met the following criteria: 1) hand speed exceeded 0.1 m/s within a 200 ms-long window before handle release (reach start) and returned below 0.1 m/s within 400 ms after handle release (reach end); and 2) no more than 4 consecutive missed frames from the camera during this sampling window. For trials that met these criteria and had 1-4 missed frames, missed marker locations were added by interpolation. Maximum speed was extracted over the window between reach start and end. Mahalanobis distance squared (MD^2^) analysis was applied to normalized hand location datasets to quantify trajectory variability (Cortes et al., 2017; Hadjiosif et al., 2021). For each trial, trajectories were first interpolated to 50 equally spaced datapoints between reach start and end. MD^2^ was calculated for x and y time series with respect to all baseline trials of the same movement direction and averaged per trial (AMD^2^). Median AMD^2^ was calculated over each week.

**Figure 2.**
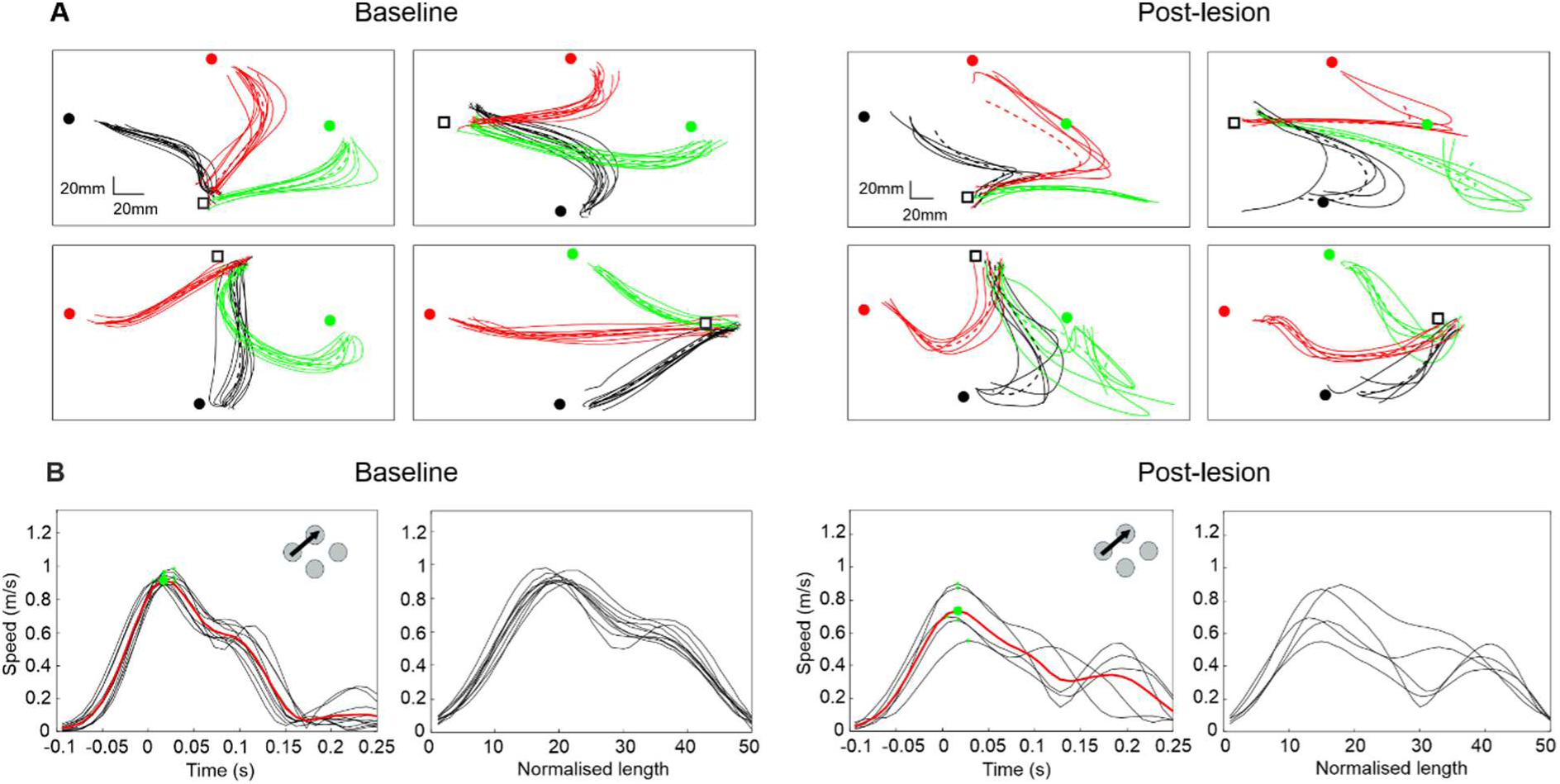
Example kinematics. A, reach trajectories plotted from DeepLabCut predictions of hand location converted to physical distance (mm). Results are from Monkey D following a New M1 and S1 Brodmann’s Area 3a/b lesion. Plots shown are taken from an example baseline session (left four plots) and early post-lesion session (right four plots). Each plot represents a different handle location (start of reach, black square), with three corresponding cup locations (end of reach; black, red or green circles). Repeats of the same reaching movement are overlaid in the same color. Dotted lines represent mean trajectories. B, hand speed plotted for all trials for one movement direction, shown from same baseline and post-lesion sessions as in 2A. The left most plot for each session represents hand speed plotted 0.1 s before to 0.25 s after handle release (time 0 s). Green dots are the maximum speeds per trial; red traces are the mean speed averaged over all trial repeats. The right most plots display overlaid speed traces, interpolated to a standardized length of 50 samples from when the hand first exceeds (reach start) and then returns below (reach end) 0.1 m/s.

Maximum speed and AMD^2^ were averaged over all movement directions to appreciate overall changes. We also found the difference between maximum speed and AMD^2^ between trials involving elbow flexion *vs* elbow extension (Fig. 1B); this allowed us to check for relative differences depending on movement type. For this difference analysis, the standard error of the median per week was estimated using bootstrapping with 1000 iterations. Significant changes in maximum reaching speed and median MD^2^ each week compared to the averaged baseline value for each monkey were identified using the two-sample t-test. Significance for both maximum speed and AMD^2^ calculations was corrected for multiple comparisons using a Benjamini-Hochberg correction with a false discovery rate of 5%.

### Magnocellular Red Nucleus Lesions

All lesion surgeries described in this report were conducted under general anesthesia managed by a veterinary surgeon. Monkeys were initially sedated with an intramuscular injection of ketamine (2-10 mg/kg), medetomidine (3 μg/kg) and midazolam (0.3 mg/kg). During surgical preparation, anesthesia was maintained with sevoflurane inhalation (2-3%). Animals were intubated and ventilated with positive pressure ventilation. A continuous infusion of alfentanil (24-108 µg/kg/hr), ketamine (6 mg/kg/hr), midazolam (0.3 mg/kg/hr), methylprednisolone (5.4 mg/kg/hr), and Hartmann’s solution (5 ml/kg/hr) ensured adequate anesthesia, analgesia, anti-inflammation and hydration throughout surgery. The animal was kept warm with a temperature-controlled heating blanket, and also a supply of thermostatically controlled warm air. Vital sign monitoring included pulse oximetry, heart rate, blood pressure, core and peripheral blood temperature, and end tidal CO_2_. Anesthetic doses were adjusted as necessary to maintain a stable plane of deep anesthesia.

In two monkeys, unilateral RNm lesions were performed using an electrocoagulation technique. Animals were placed in a stereotaxic frame; coordinates in the following are given relative to inter-aural line at the midline in anterio-posterior (AP), mediolateral (ML) and dorso-ventral (DV) directions. A craniotomy was made on the left side stretching from AP 2-12A and ML 3.5-6.5 (Martin and Bowden, 1996; Belhaj-Saïf et al., 1998). We began by mapping the oculomotor nuclei, as these are close to RNm and produce easily visible eye movements upon stimulation. A stainless-steel electrode (MS501G, Micro-Probe, San Jose, US) penetrated at a 5° penetration angle, and was initially positioned to target coordinates AP 7.0A ML 0 DV 5.0. A series of penetrations at various heights and AP locations close to these initial coordinates were made, and single pulse stimuli (biphasic, 0.1 ms per phase) were delivered at each site. Where eye movements were elicited, the intensity was reduced to find the threshold. These were noted on graph paper, producing a fine-grained map of the oculomotor nuclei. The electrode was then repositioned to target an initial site 1 mm to the left of the optimal oculomotor nucleus coordinate, and RNm mapping begun.

The RNm should facilitate motoneurons via rubrospinal pathways; however, under general anesthesia motoneuron excitability is reduced, and it can be difficult to elicit overt activity by stimulating weak inputs. To overcome this, we first delivered a train of stimuli (biphasic pulses, 0.1ms per phase, 700-1000 µA intensity, 3 ms inter-stimulus interval) to the pyramidal tract, and checked for responses in electromyogram (EMG) from the right extensor digitorum communis (EDC) muscle. Pyramidal tract stimulation used a chronic electrode implanted in a previous surgery (not otherwise used in the work reported here). The number of stimuli in the train was increased progressively until an EMG response was produced and was then reduced by one to obtain a just subthreshold stimulus. The mapping electrode was then advanced to various sites surrounding the expected location of RNm, and stimuli given through it following the subthreshold pyramidal tract stimulation (0.4 ms interval between the last stimulus in the train to the pyramidal tract and the RNm stimulus). We searched for locations where EDC responses were elicited and determined the threshold intensity of RNm stimulation required (25 µA-1 mA). These thresholds were plotted on graph paper, as for the oculomotor nuclei. Lesion locations were determined as the lowest threshold sites. Electrophysiological recordings from the right EDC muscle during mapping at the chosen lesion sites for each monkey are presented in Supplementary Figure 1.

A direct current was passed at each chosen site to produce a lesion (1 mA for 20 s, or 2 mA for 10 s, electrode positive). Monkey Cm had two lesions at the following stereotaxic locations: AP 6.0A, ML 1.0L, DV 16.9 and AP 7.0A, ML 1.0L, DV 18.0. Monkey Ca had three lesions at AP 5.0A, ML 1.0L, DV 14.6; AP 6.0A, 1.0L, DV 14.3 and AP 7.0A, ML 1.0L, DV 13.7. The electrode used for mapping and lesioning was then removed, and the craniotomy sealed. Both animals recovered from anesthesia and returned to their normal cages later that evening. Broad spectrum antibiotics (oral or subcutaneous co-amoxyclav 12.5 mg/kg), analgesics (intramuscular buprenorphine 0.02 mg/kg; oral or intramuscular meloxicam 0.2 mg/kg; oral paracetamol 23 mg/kg) and steroids (intramuscular dexamethasone 0.2-0.5 mg/kg) were administered. Two days later both animals were well enough to return to the laboratory to perform the task. They received a secondary cortical lesion when kinematic measures plateaued after 7-8 weeks (Monkey Cm – Anterior Old M1; Monkey Ca – Entire Old M1).

### Cortical Lesions

Monkeys were placed in a stereotaxic frame under general anesthesia (details as above) and a craniotomy made on the left side above M1 (approximately 6-20 mm lateral to midline). This lesion laterality was chosen to cover much of the right upper limb representation (8-19 mm) (Boudrias et al., 2010). The anteroposterior extent varied depending on the animal and the intended lesion.

To make lesions of different M1 subregions at the precentral gyrus (i.e. Old M1), the dura was opened to allow for smooth needle penetration. A manipulator was configured with four needles oriented anteroposterior, spaced 1 mm apart, and pre-filled with 900 µM endothelin-1 in saline (HB3442, Hello Bio, Bristol, UK). Endothelin-based approaches for generating focal ischemic cortical lesions have already been used effectively in macaques (Herbert et al., 2015), due to endothelin’s fast-acting and potent vasoconstrictive effects. We anticipated a diffusion radius of approximately 0.5 mm relative to the needle tip. Anteroposterior positioning of needles relative to the central sulcus (CS) determined which lesion was performed (Posterior Old M1 1-4 mm anterior, blue in Supplementary Fig. 2A; Anterior Old M1 4-7 mm anterior, purple). Starting 8mm lateral from the midline, needles first penetrated 2 mm below the surface and were then retracted to 1.3 mm below the surface; this allowed for some tissue dimpling on penetration. Endothelin-1 was infused over two minutes (1 μl per site, 500 nl/min) with needles left in the tissue for a further minute to allow sufficient diffusion. Needles were then moved to the next site 1 mm more lateral. The injections were repeated identically at least 1 hour later to increase the duration of ischemia (Herbert et al., 2015). In one animal (Monkey Ca), a larger lesion was performed 7 weeks after the RNm lesion. The same method was used but with an increased number of anteroposterior infusion sites (Entire Old M1, 1-7 mm anterior relative to the CS).

To perform deeper lesions of the grey matter in the anterior bank of the CS (New M1), we first used electrophysiological mapping to determine the angle of the CS. A parylene-insulated stainless steel microelectrode (WE30031.0A3, MicroProbe, San Jose, USA) was zeroed mediolaterally to stereotaxic midline and anteroposteriorly to the CS. A ground wire was connected to a needle placed in the skin. The electrode penetrated into the cortex, and single unit recordings were made, noting whether cells were discharging at depths of 0.5-10 mm below the surface. This was repeated for anteroposterior sites 0.5-4 mm anterior to the CS, and at 9.5, 13.5 and 17.5 lateral to midline. The anteroposterior sites at each laterality that had the largest and most consistent cell firing were used as lesion targets, as these were likely to be within cortical layer V. A four-needle array was configured in a manipulator with 1 mm spacing and oriented approximately mediolaterally, with the angle adjusted to ensure the needles were parallel to the CS. The needles were penetrated to 6.7 mm below surface before being retracted 0.7 mm to allow for tissue dimpling. Endothelin-1 was infused at each site as above. The needles were moved 1 mm higher, and the infusion process repeated to cover 3-6 mm below cortical surface (orange in Supplementary Fig. 2A). Three such penetrations were made at different medio-lateral locations to cover 8-19 mm from the midline.

At the end of these lesion surgeries, the dura was sutured and craniotomy sealed using Gelfoam (Pfizer, Kent, UK) and dental acrylic (Kemdent, Swindon, UK). Animals recovered in a padded cage under supervision. They received a full program of postoperative care (oral/subcutaneous co-amoxyclav 12.5 mg/kg or oral doxycycline 8 mg/kg; oral/intramuscular meloxicam 0.2 mg/kg; oral paracetamol 23 mg/kg; intramuscular dexamethasone 0.2 mg/ml). All monkeys were well enough to return to their home cages the morning after and returned to the laboratory for recording 2 days post-lesion.

We performed a more extensive cortical lesion in one animal (Monkey D), targeting PMd, Entire M1 and S1 (Brodmann’s Areas 1, 2 and 3). This used a combination of the superficial and deep lesioning approaches described above, split into three surgeries each separated by one week. The first surgery entailed mapping both anterior and posterior banks of the CS (New M1 and S1 Brodmann’s Area 3a&b, respectively). A craniotomy was made extending 13 mm anterior and 9 mm posterior relative to the CS to cover the final desired lesion extent. The dura was left intact. Maps of cell discharge were compiled from 3 mm anterior to 1 mm posterior relative to the CS, at 9, 12, 15 and 18 mm lateral to the midline, and between 1-12 mm below the cortical surface. The craniotomy was then sealed as above.

One week later, the dental acrylic over the craniotomy was removed, and a surgery to produce deep lesions of anterior and posterior banks of the CS was carried out. Two manipulators holding either a four or six-needle array were pre-filled with endothelin-1 and oriented antero-posteriorly with 1 mm spacing. The most posterior needle of each array was zeroed to midline. The four needle array sites were penetrated in turn from 8-13 mm lateral, the six needle array from 14-19 mm lateral. At each laterality in turn, the needles penetrated to an appropriate depth and anteroposterior location relative to the CS as determined from mapping to cover both pre- and post-central sulcus. Endothelin-1 infusions were made over 1.5 minutes (1 μl per site, 667 nl/min). Needles were left in place for a further minute before being moved to the next site 1 mm higher. When all depths were lesioned (i.e. from initial infusion depth to 2 mm below surface, pink in Supplementary Fig. 2B), the needle arrays were withdrawn, moved 1 mm more lateral, and the protocol repeated. All lesions were repeated after approximately one hour as described above. At the end of surgery, the craniotomy was sealed and appropriate post-operative care given (both as previously described). Monkey D was well enough to return to the laboratory and the task two days after this first lesion.

A final surgery to lesion superficial cortical regions both anterior and posterior to the CS (PMd, Old M1, S1 Brodmann’s Areas 1 and 2) was performed one week later. The protocol used was similar to that previously described for lesions of Old M1, except that two needle arrays with four or six needles oriented antero-posteriorly were used simultaneously to cover a larger anteroposterior extent (green in Supplementary Fig. 2B). In a first set of injections, the six needle array covered 1-6 mm anterior to CS, and the four needle array 1-4 mm posterior to CS. These injections were repeated after one hour. In the second set, injections covered 7-12 mm anterior and 5-8 mm posterior to CS, respectively; these were also repeated after one hour. At the end of the surgery the dura was repaired, the craniotomy sealed, and appropriate post-operative therapy given (as preciously described). Laboratory recording restarted two days after this second lesion.

### Cortical Histology

At 15 weeks post-lesion, animals were terminally anesthetized with an overdose of propofol and perfused through the heart with phosphate buffered saline, followed by formalin for fixation of tissues. The brain and brainstem were removed and immersed in formalin for 24 hours, then progressively submerged in ascending concentrations of sucrose solution (10%, 20%, 30%) for cryoprotection.

Post-mortem histological analysis was performed to verify the extent of cortical lesions in each animal. The left and right sensorimotor cortices were dissected from midline to approximately 21 mm lateral, and a cryostat used to cut 50 µm thick parasagittal sections from each hemisphere. Sections were mounted every millimeter from 8-19 mm to cover the upper limb representation. For some animals, sections ranging the full mediolateral extent were not available due to technical limitations. In these cases, analyses were restricted to the subset to which sections were available (Monkey Cu: 8-18 mm; Monkey Co: 11-19 mm; Monkey Zd: 8-16 mm; Monkey Ze: 9-14 mm).

For Monkeys Zd and Ze, an immunohistochemistry protocol using fluorescently labelled antibodies for glial fibrillary acidic protein (GFAP; donkey anti-rabbit IgG, Alexa Fluor 488; Thermo Fisher Scientific, Massachusetts, USA) and neuronal nuclear protein (NeuN; donkey anti-mouse IgG, Alexa Fluor 488; Thermo Fisher Scientific, Massachusetts, USA) were used to visualize chronic glial scarring and neuron-specific cell loss following the lesions, respectively. For Monkeys B, Ca, Ch, Cm, Co, Cu, D, available sections were stained with cresyl violet, which allowed for visualization of gross cytomorphology.

Brightfield images of stained sections were analyzed using OMERO Plus Software (v5.29.2, (Allan et al., 2012)). Firstly, lesion outlines were drawn manually on each section on the lesioned (left) side, defined following close visual inspection of cytomorphological abnormalities. When outlining sections with no surviving cortical surface (example in Supplementary Fig. 3A), non-lesioned right hand side tissue at the same mediolateral extent was used as a reference of healthy anatomy. Next, the anteroposterior (AP) boundaries of each cortical region were drawn on each image, based on physical distances defined in a recent mapping study in monkeys (Boudrias et al., 2010); these boundaries were (from most anterior to posterior): PMd = 7-12 mm anterior to CS (5 mm AP extent); Anterior Old M1 = 4-7 mm anterior to CS (3 mm AP extent); Posterior Old M1 = 1-4 mm anterior to CS (3 mm AP extent); New M1 (superior extent) = 1mm inferior from layer V in Posterior Old M1; New M1/S1 Area 3 (inferior extent) = most inferior, large layer V cells in bank of CS, 1 mm anterior from start of CS; S1 Area 3 superior extent = 1 mm inferior from layer V in S1 Areas 1&2, 1mm posterior from start of CS; S1 Areas 1&2 = 1-8 mm posterior of the CS posterior bank (7 mm AP extent). Cortical region boundaries are illustrated in Supplementary Figure 3B. The distance of layer V cells affected by the lesion in each cortical region was drawn using straight lines, and physical distance in millimeters extracted. This process was repeated with sections spaced 1 mm apart from each other in the mediolateral direction, to cover the full extent of the lesion injections.

The proportion of layer V tissue damaged was quantified per section *(i)* and per cortical region (*P*_damage_); *D*_damaged_ is the anteroposterior distance (mm) over which layer V neurons were damaged, and *D*_total_ is the total anteroposterior extent (mm) of the cortical region.

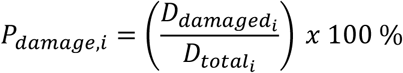

*P*_damage_ values were rounded to the nearest whole numbers for Monkeys B, Ca, Ch, Cm, Cu, Co D. For Monkeys Zd and Ze, *P*_damage_ values were assigned into either 0%, 25%, 50%, 75% or 100% bins; this was due to an alternative staining methodology used for these monkeys, which resulted in lower confidence predictions of lesioned layer V compared to the cresyl violet approach.

The overall percentage of layer V cells damaged in each cortical region across the available mediolateral extent (*P*_damage, overall_) was calculated using an average of *P*_damage_ values over the number of sections stained (*n*). This process was repeated for each monkey.

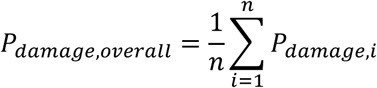

The full analysis pipeline was conducted separately by two of the authors independently (AB and AMEB) and any discrepancies resolved, to reduce bias and increase accuracy. Results from this analysis are shown in Table 1 and Figure 3.

**Figure 3.**
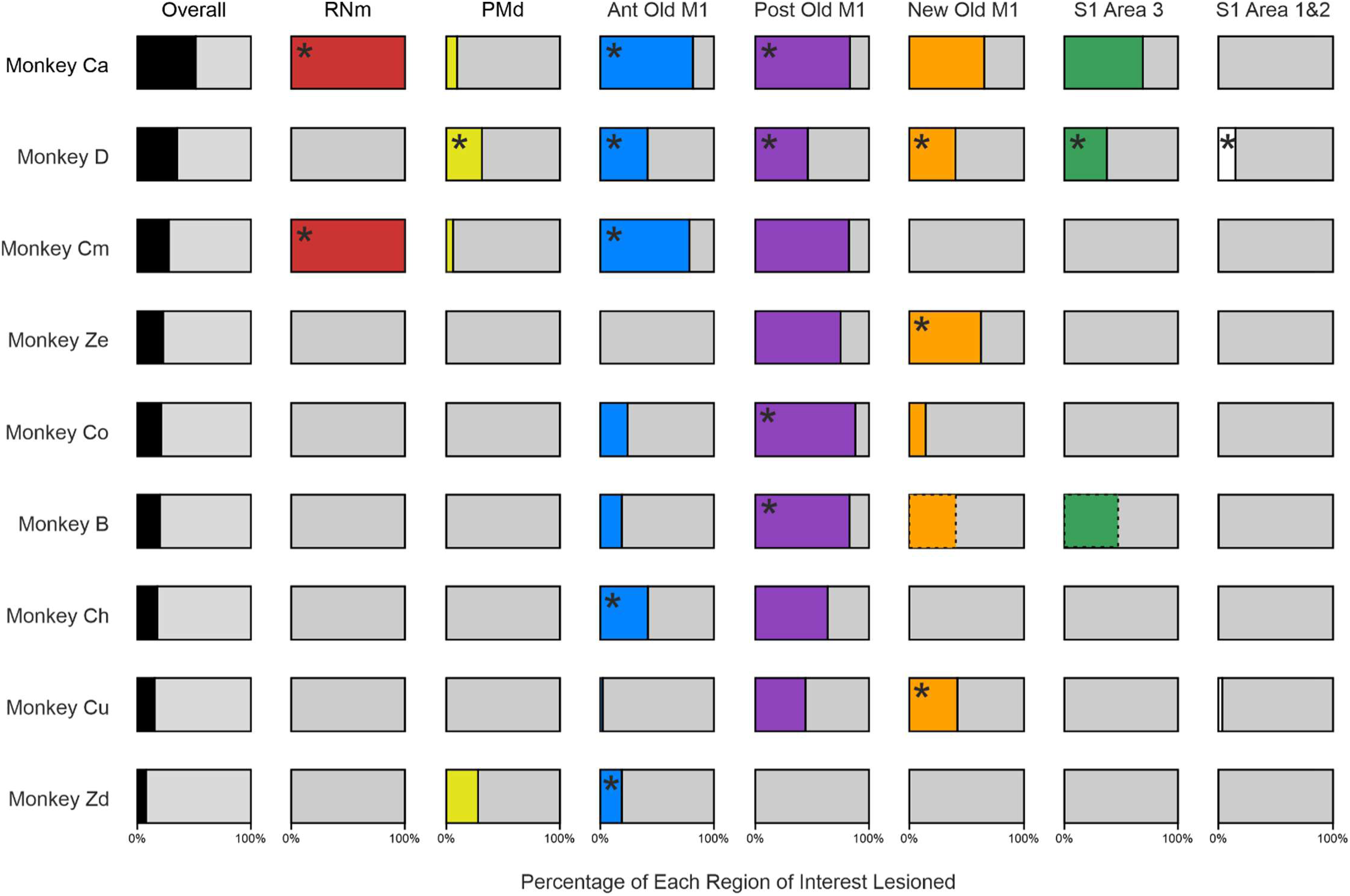
Visual representation of results from Table 1. Regions marked with an asterisk are the lesion targets for that animal. Overall values were calculated using an average of damage across all cortical regions in each monkey. RNm results have been expressed as 100% or 0% as we do not have numerical values for this lesion type. Bars representing damage in New M1 (orange) and S1 Area 3 (green) for Monkey B are presented with dotted lines; these represent the more medial values calculated from 3.5-14.5 mm from midline in this animal due to considerable damage noted to both banks of the sulcus in this monkey only.

**Table 1.** Summary of regions damaged from lesions in each monkey. Data were obtained from post-mortem histological analysis. Rows separate monkeys, ordered from greatest overall damage to cortical layer V cells (Monkey Ca) to least (Monkey Zd). Columns separate cortical regions; numbers represent the percentage of layer V cells in that cortical region that were affected by the lesion. No numerical data were available for RNm lesion verification – the analysis was qualitative. * denotes percentage damage calculated from a more medial extent (3.5-14.5 mm from midline), where there was substantial damage to both anterior and posterior banks of the sulcus for Monkey B, not seen in the other animals.

| Monkey | RNm Lesion? | Average Across Cortical Regions | PMd | Anterior Old M1 | Posterior Old M1 | New M1 | S1 Area 3 | S1 Areas 1&2 |
| --- | --- | --- | --- | --- | --- | --- | --- | --- |
| Monkey Ca | Yes | 51 | 10 | 82 | 83 | 65 | 69 | 0 |
| Monkey D | - | 35 | 31 | 42 | 46 | 40 | 37 | 15 |
| Monkey Cm | Yes | 28 | 6 | 78 | 82 | 0 | 0 | 0 |
| Monkey Ze | - | 23 | 0 | 0 | 75 | 63 | 0 | 0 |
| Monkey Co | - | 21 | 0 | 24 | 88 | 14 | 0 | 0 |
| Monkey B | - | 20 | 0 | 19 | 83 | 7 (40*) | 10 (49*) | 0 |
| Monkey Ch | - | 18 | 0 | 42 | 64 | 0 | 0 | 0 |
| Monkey Cu | - | 15 | 0 | 2 | 44 | 42 | 0 | 3 |
| Monkey Zd | - | 8 | 28 | 19 | 0 | 0 | 0 | 0 |

### Midbrain Histology

We performed post-mortem histological analysis using midbrain tissue from Monkeys Cm and Ca, which allowed localization of the electrolytic lesion sites. Cryoprotected midbrain was dissected from the brainstem, and 50 µM thick coronal sections were cut using a cryostat. Sections from both animals were stained with cresyl violet to visualize cytoarchitecture. In Monkey Ca, an additional Perls Prussian Blue protocol with Nuclear Fast Red counterstain was used to help verify lesion locations. This method produced a blue precipitate at sites containing ferric iron deposits left behind by the stainless-steel lesion electrode. Brightfield images of all stained sections were visually inspected using OMERO Plus Software (v5.29.2, (Allan et al., 2012)). Multiple lesion sites were located in each animal (Fig. 4).

**Figure 4.**
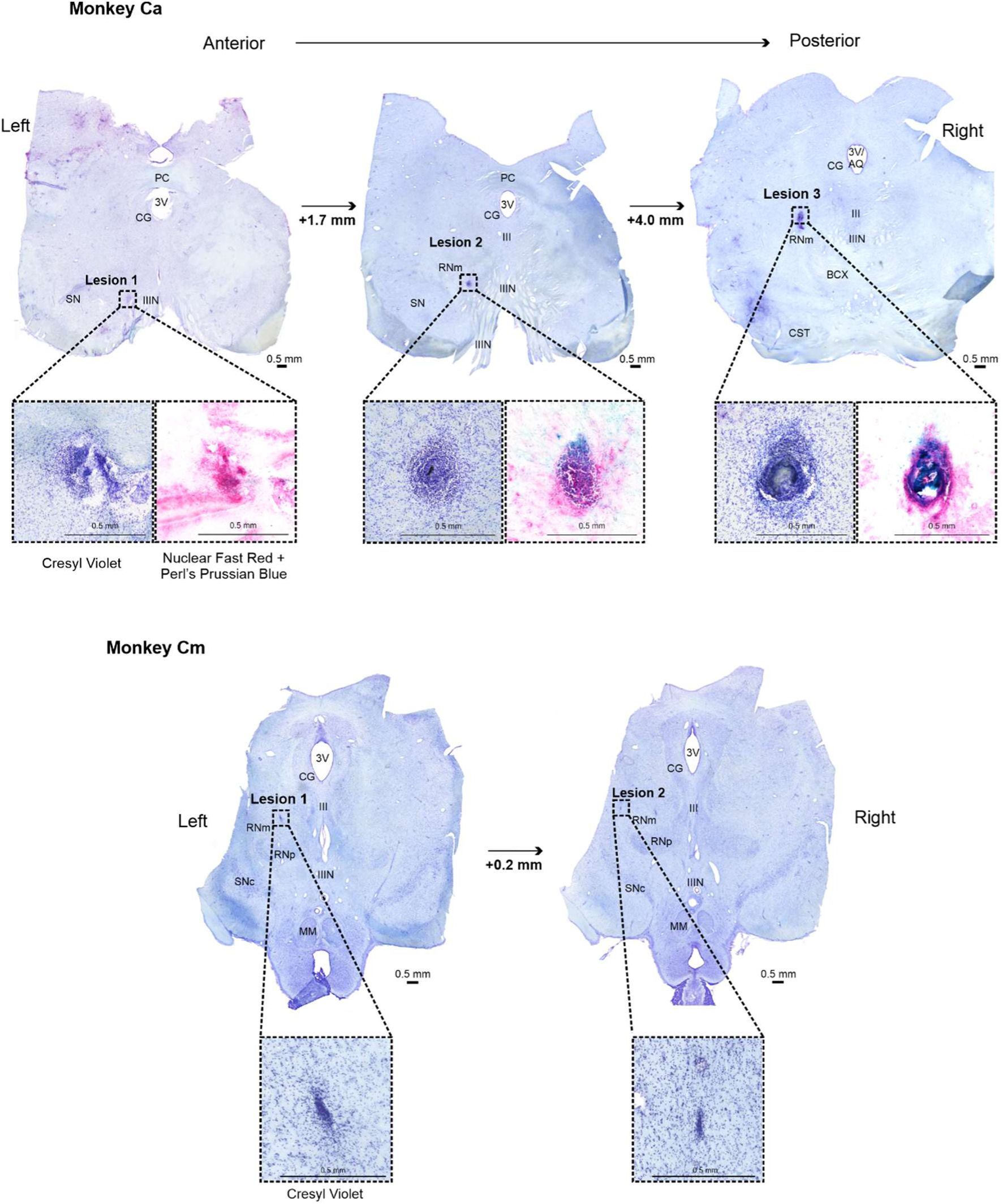
RNm lesion locations confirmed with histological staining for Monkeys Cm and Ca. 50 µm-thick midbrain sections from both monkeys were stained with cresyl violet to visualize Nissl bodies. Additional counterstaining in alternate sections was conducted for Monkey Ca using Nuclear Fast Red and Perl’s Prussian Blue, allowing for visualization of iron deposits left behind by the lesion electrode. Sections were photographed under bright field illumination. Each column shows an individual lesion site at its largest extent (three in Monkey Ca; two lesions in Monkey Cm). High resolution images of each lesion site are shown next to the corresponding whole midbrain section. *AQ = aqueduct; BCX = decussation of the brachium conjunctivum; CG = central gray; CST = corticospinal tract; MM = mammillary bodies; PC = posterior commissure; RNm = magnocellular red nucleus; RNp = parvocellular red nucleus; SN = substantia nigra; SNc = substantia nigra pars compacta; IIIN = oculomotor nerve; III = oculomotor nucleus; 3V = third ventricle*

## Results

### Extent of Cortical Lesions

The aim of this study was to conduct selective lesions of sensorimotor cortical regions and assess the kinematic impacts on reaching in monkeys. Table 1 displays data from post-mortem histological analysis, arranged in rows for each monkey, and columns of percentage lesion damage to cortical layer V, overall and in each cortical region. Figure 3 summarizes Table 1 results in a visual format; asterisks mark lesion targets in each animal. Monkeys are organized from greatest overall cortical lesion damage (Monkey Ca) to the least (Monkey Zd). These visual bar charts are referenced in later kinematics figures for comparison between the cortical areas lesioned and corresponding kinematic effects.

In all animals, we successfully damaged the target cortical regions (Monkey Ca – Entire Old M1; Monkey D – All regions of interest; Monkey Cm, Ch and Zd – Anterior Old M1; Monkeys B and Co – Posterior Old M1; Monkey Ze and Cu – New M1), albeit with varying severity.

No animals had lesions isolated to a single cortical area; damage always spread across multiple regions. Monkey Ca had the largest degree of spread, with substantial collateral damage to New M1 and S1 Area 3, despite the lesion intended to encompass only Old M1. A surprising finding was that 8/9 animals had considerable damage to Posterior Old M1, regardless of the lesion target.

In addition, lesions to a target cortical region were never complete; all animals retained some portion of the target region’s upper-limb representation. This was most apparent in Monkey Zd, where damage to the target region was substantially less extensive than anticipated, with involvement of 19% layer V cells.

These findings collectively indicate that although endothelin-1 injections consistently involved the intended cortical region, the efficacy and extent of unintended damage varied across animals. Consequently, all results in this study will henceforth be presented according to the actual lesion locations determined from histological analysis rather than the intended targets.

### Extent of RNm Lesions

In these experiments, we intended to lesion the left RNm selectively in two animals (Monkey Cm and Ca). Figure 4 shows post-mortem histological analysis of coronal midbrain sections which were used to confirm the locations of electrolytic lesions (Monkey Cm -top two images; Monkey Ca -bottom three images). Sections are presented from most anterior (left) to posterior (right). All tissue was stained with cresyl violet, wherein abnormal cellular aggregation served as the histological marker of lesioned tissue. Each section has been annotated with anatomical landmarks(Mikula et al., 2007) and contains a different electrolytic lesion (emphasized with black dotted square). Each lesion site is also shown at a higher magnification for clarity (larger black dotted boxes). For Monkey Ca, there are additional high magnification images of lesions stained blue, indicative of iron deposits left behind by the lesioning electrode.

Two out of three lesions identified in Monkey Ca were located within the RNm. Lesion 3 (the largest) was positioned approximately 2mm to left of the oculomotor nucleus (III) and the oculomotor nerve fibers (IIIN). The decussation of the brachium conjunctivum (BCX) was visible inferiorly. This lesion site included the RNm (Mikula et al., 2007), although it may have extended to involve the neighboring superior cerebellar peduncle (SCP) (Martin and Bowden, 1996). We observed no clear signs of ataxia or ocular-motor abnormalities in this animal, so any SCP involvement was likely to be minimal.

Lesion 2 was located approximately 3.9 mm more anterior and 2mm inferior relative to lesion 1. The BCX was no longer present. The oculomotor nucleus and axons (IIIN and III) were present, along with the substantia nigra pars compacta (SNc) more inferiorly. We would expect the RNm to be approximately 1-2 mm lateral from the IIIN (Martin and Bowden, 1996; Mikula et al., 2007); this is concordant with the position of lesion 2. It is worth noting that the magnocellular and parvocellular (RNp) divisions of the red nucleus have close spatial proximity (RNm is more lateral than RNp), which makes it very difficult to lesion the RNm selectively. It is possible damage could have extended to RNp, which may have contributed to post-lesion reaching deficits in Monkey Ca, similar to such reported in rodents with selective RNp lesions (Morris et al., 2015).

Lesion 1 was the least severe and least accurately placed of the three lesions in Monkey Ca. It was located approximately 1.9 mm more anterior relative to lesion 2 and appears too inferior (nearby/within the IIIN). At no point during recovery did we observe any ocular motor signs (for example, ptosis, pupillary abnormalities). Because such signs would be highly conspicuous following genuine IIIN damage, lesion 1 most likely produced no meaningful oculomotor nerve injury.

Both electrolytic lesions identified in Monkey Cm were less severe in comparison to Monkey Ca, and were located closely relative to each other (within 0.2 mm in the anteroposterior direction). The RNp and SNc are visible in both sections (similar anatomy to Monkey Ca, lesion 2). Lesion 1 in Monkey Cm was found approximately 1 mm more superior than RNp at a similar laterality (approximately 1.5 mm from midline). Lesion 2 was at the same depth as lesion 1, but approximately 1 mm more lateral. Both of these lesions were within the RNm.

Overall, these results indicate successful targeting of the RNm in both animals, although the lesions appeared to be larger and likely more severe in Monkey Ca.

### Impact of Cortical Lesions on Reaching Performance

None of the lesions reported here resulted in an overt clinical presentation resembling the flexor synergy seen in human stroke survivors -that is, elbow flexion occurring in conjunction with shoulder abduction. Application of sensitive quantitative measures capable of detecting flexor synergy in a subset of these animals will be discussed in a future publication. In this report, we focus on differences in negative motor phenotypes.

Figure 2 shows example kinematic measurements before and after a lesion. In Figure 2A, reach trajectories are shown for each of the 12 possible movements, from the handle starting position (white square) to the cup (three locations shown in red, green and black on each plot). Trajectories post-lesion were clearly more variable. Figure 2B illustrates the instantaneous speed plots for a single movement direction. The left trace shows raw data, with an x-axis in seconds; each movement lasts a slightly different time. The right traces are normalized in length between movement onset and offset; data processed in this way were used for the AMD^2^ analysis. Movements after the lesion were a little slower, more variable, and in some cases had pronounced two components.

We present changes to two kinematic assessments pre- and post-lesion for each monkey: AMD^2^ (Fig. 5) and maximum speed (Fig. 6). These measures were averaged over all reaching directions to assess overall reaching performance regardless of any differences in reaches requiring elbow flexion or elbow extension. AMD^2^ provides a quantitative measure of trajectory variability. Increased AMD^2^ scores reflect a reduced ability to control muscles independently which is required for accurate straight line reaching movements and may thus be a proximal analogue of lowered dexterity in the hand. Reduced maximum speed by contrast may indicate weakness. In all monkeys scores after the lesion were compared with the pre-lesion baseline. Most recovery of both kinematic measures occurred 1-4 weeks post-lesion (the spontaneous recovery period), similar to the acute/subacute recovery window previously described in humans (Cortes et al., 2017). Recovery usually plateaued within ∼8 weeks, similar to humans (van Kordelaar et al., 2014; Cortes et al., 2017).

**Figure 5.**
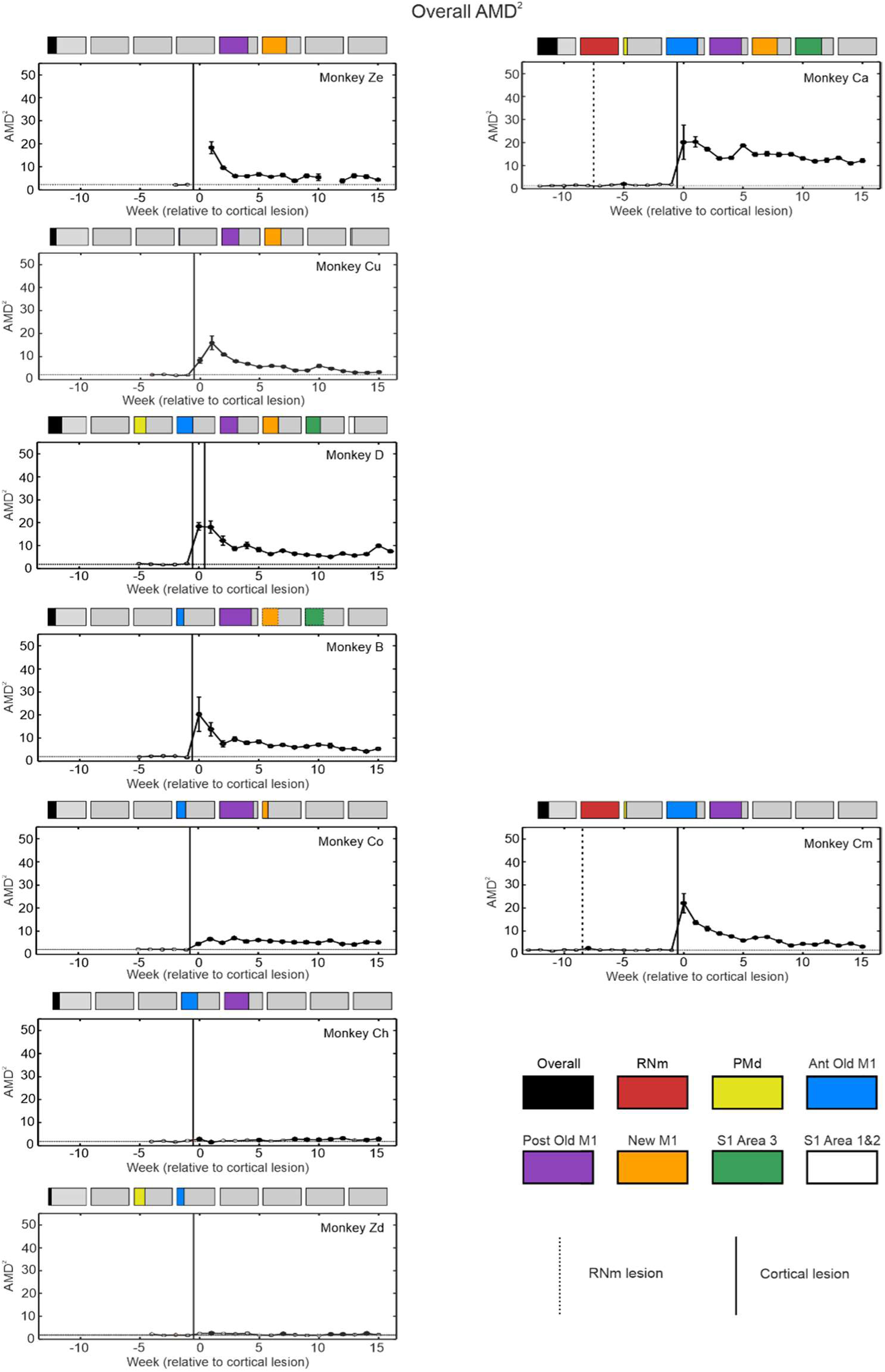
Trajectory variability per week averaged over all reaching directions. Each graph contains average Mahalanobis distance squared values (AMD^2^) per week for each monkey. Higher AMD^2^ scores represent poorer reaching quality. Coloured bars above each plot show the percentage of each cortical region lesioned. Plots have been arranged in an order of largest degree of New M1 damage (orange bar) at the top, to the smallest at the bottom. Vertical lines show lesion times (RNm or cortical); horizontal lines indicate baseline mean ± standard error; circles are median values from all trials performed that week. Standard error of the median was calculated by bootstrapping with 1000 iterations (error bars). AMD^2^ ranges on the y axis are the same across all plots to allow easier visual comparison. Filled circles are significantly different from baseline (p < 0.05), identified using the two-sample t test and corrected for multiple comparisons using the Benjamini-Hochberg procedure with a false discovery rate of 5%.

**Figure 6.**
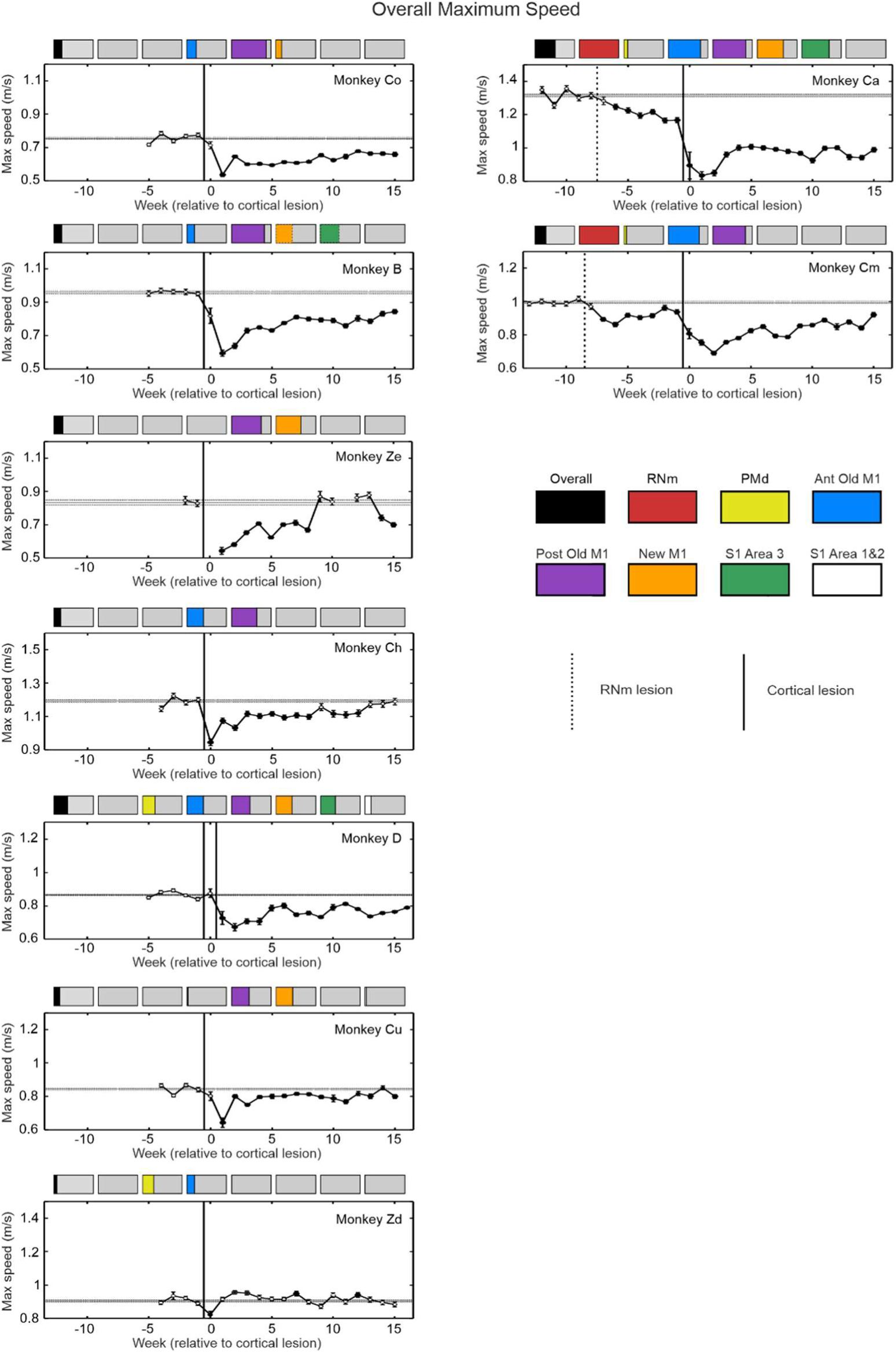
Maximum reaching speed per week averaged over all reaching directions. Each graph contains average maximum speed per week from an individual monkey. The coloured bars above each plot show the percentage of each cortical region lesioned. Plots have been arranged in an order from largest degree of Posterior Old M1 damage (purple bar) at the top, to the smallest at the bottom. Vertical lines show lesion times (RNm or cortical); horizontal lines indicate baseline mean ± standard error; circles represent mean weekly values, error bars show standard error of the mean. Filled circles are significantly different from baseline (p < 0.05), identified using the two-sample t test and corrected for multiple comparisons using the Benjamini-Hochberg procedure with a false discovery rate of 5%.

Comparing cortical lesions with varying degrees of involvement across the different divisions of M1 (New M1 vs Posterior Old M1 vs Anterior Old M1) revealed different initial deficits and subsequent recovery outcomes. Figure 5 plots are arranged in an approximate order of New M1 damage (orange bar), with the greatest involvement displayed at the top (Monkeys Ze and Ca). All monkeys with lesions that affected New M1 had significant negative impacts to trajectory variability (Monkeys Ze, Ca, D, Cu, B, Co), irrespective of severity. Post-lesion AMD^2^ values remained permanently higher than baseline in all these animals. In contrast, monkeys with lesions that spared New M1 (Monkeys Ch and Zd) showed only minimal, if any, increases to AMD^2^. The sole exception was Monkey Cm, who displayed large and significant subacute impairment to trajectory variability despite sparing of New M1; this could be attributable to the preceding RNm lesion. Damage to the RNm may also explain why Monkey Ca had the poorest recovery of this measure, with a combined RNm and New M1 lesion.

Plots of maximum speed (Fig. 6) are organized according to the degree of damage to Posterior Old M1 (purple bar). All lesions affecting Posterior Old M1 caused significant slowing of maximum reaching speed, which was most sustained in monkeys with the largest percentage damage in this region (Monkeys Ze, Ca, B, Cm, Co). Monkey Zd had no damage in Posterior Old M1, and reaching speed was barely affected. Previous work has shown that Posterior Old M1 has mixed outputs via both corticospinal and corticoreticulospinal tracts (Keizer and Kuypers, 1989), and that the majority of drive to motoneurons producing movement in primates comes from the reticulospinal tract (Tapia et al., 2022). We suggest that without strong facilitation of motoneurons via the cortico-reticulospinal tract, permanently slower movements resulted after Posterior Old M1 lesions.

Conversely, Anterior Old M1 lesions did not appear to exert additive negative effects on our kinematic measures. In Figure 5, increasing damage to Anterior Old M1 was actually associated with progressively smaller overall effects on trajectory variability, as evidenced by the progression from Monkey B to the more extensive lesions in Monkey Co and subsequently Monkey Ch. In Figure 6, this same claim can be applied to maximum speed change: greater involvement of Anterior Old M1 did not correspond to larger overall reductions in reaching speed. In Monkey B, Co and Ch, progressively greater Anterior Old M1 damage did not result in greater speed impairments. These results suggest that New M1 and Posterior Old M1 areas have facilitatory effects on the upper limb, whereas Anterior Old M1 likely plays a different role.

In Monkey D, we tested the effect of a widespread cortical lesion (PMd, M1, S1 Brodmann’s Areas 1, 2 and 3). The effects were not as severe as anticipated, given that the only major motor cortical region left intact was the supplementary motor area (SMA). Following New M1 and S1 Brodmann’s Area 3 lesions (lesion surgery 1, first solid vertical line in Figs. 5 and 6), there was a large increase in AMD^2^ score (Fig. 5), but no change in maximum speed (Fig. 6). Subsequent lesions to PMd, Old M1 and Broadman’s Areas 1 and 2 (lesion surgery 2) had the opposite effect, with decreased speed but no further negative impact on AMD^2^. Both metrics remained significantly worse than baseline by 15 weeks post-lesion.

Interestingly, no additional deficits in quality of reaching (AMD^2^) were observed in Monkey D following combined lesions to S1 Brodmann’s Areas 3 and New M1 (Fig. 5), compared to monkeys with similar sized damage to New M1 but intact S1 Area 3 (Fig. 5 -Monkeys Ze and Cu). A similar pattern was noted in the same animal regarding reach speed (Fig. 6), with no further reduction observed after combined lesions to PMd, S1 Brodmann’s Areas 1 and 2, and Old M1, relative to monkeys with significant damage primarily isolated to Old M1 (Monkeys B and Co). These findings suggest that additional damage to somatosensory or premotor areas or New M1 did not exacerbate reach impairment beyond that caused by M1 injury alone.

### Impact of RNm followed by Cortical Lesions on Overall Reaching Performance

Two animals underwent RNm lesions (Monkeys Cm and Ca), with consistent effects (region between dotted and solid vertical lines in Figs. 5 and 6). In both there were significant and persistent reductions in speed after the RNm lesion alone (Fig. 6) but not AMD^2^ (Fig. 5), however these were smaller in amplitude compared to New M1/Posterior Old M1 lesions.

Comparing data from Monkey Cm (RNm plus Old M1 lesion) and Monkeys Co and Ch (similar size Old M1 lesions but without RNm lesion) provided insights into the contribution of the rubrospinal tract to recovery following cortical injury. The Old M1 lesion had greater initial effects on AMD^2^ after the preceding RNm lesion than when the cortical lesion occurred against the background of a healthy rubrospinal outputs. The deficits persisted until the end of the recorded period.

In Monkey Ca, an entire M1 and S1 Area 3 lesion after the preceding RNm lesion led to the poorest recovery seen in our study, with large deficits in both AMD^2^ and speed which improved little. Notably, this animal was more severely affected than Monkey D, who had lesion damage in these same cortical regions but without a preceding RNm lesion (albeit the extent of damage of each area was less). These results therefore serve to highlight further the importance of RNm as a route for recovery in monkey.

Overall, the results from RNm lesioned animals suggest a role for the rubrospinal system in generating fast reaches, and in contributing to recovery after cortical damage. Also, the data further emphasize the detrimental consequences of losing New M1 & S1 Brodmann’s Area 3 in addition to Old M1, as this is a considerable difference separating Monkey Cm (RNm plus Old M1, better overall recovery) and Monkey Ca (RNm plus entire M1 & S1 Brodmann’s Area 3, much poorer overall recovery).

### Impact of Cortical Lesions on Reaching Requiring Elbow Flexion vs Elbow Extension

Imbalanced recovery is a common finding in stroke survivors, who are often left with overactive flexors and weak extensors (Kamper et al., 2003). We examined whether there was a difference in impairment between movements which required elbow flexion or extension, by plotting differences between AMD^2^ (Fig. 7) and maximum speed (Fig. 8) for flexion vs extension.

**Figure 7.**
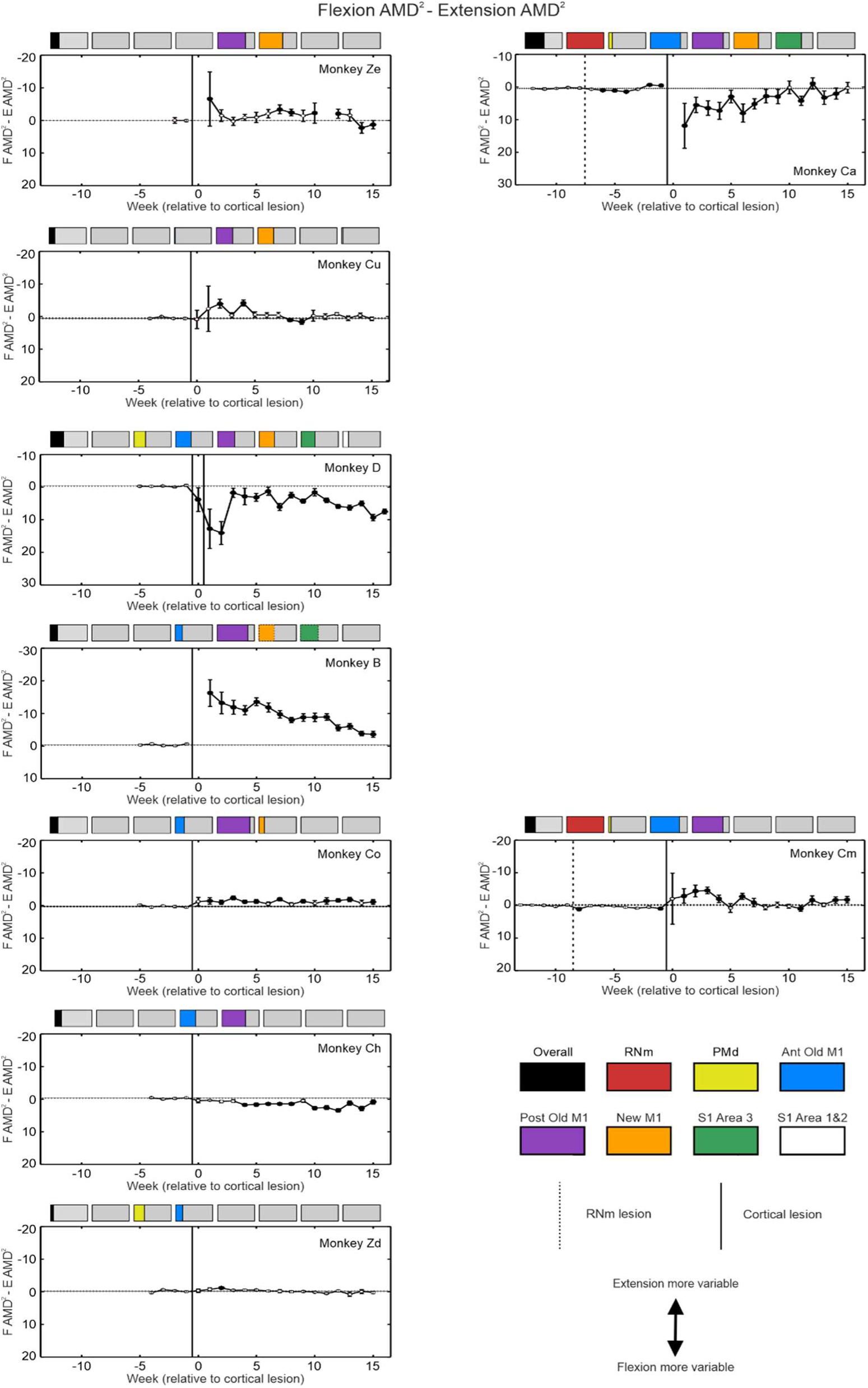
Maximum trajectory variability per week compared for elbow flexion vs extension reaching directions. Similar display as Fig. 8, but results shown are the difference between flexion and extension values. The y axis range is comparable across all plots between monkeys to allow easier visual comparison. Note that the axis scale has been inverted, so that worse performance in extension is shown by upwards deflection.

**Figure 8.**
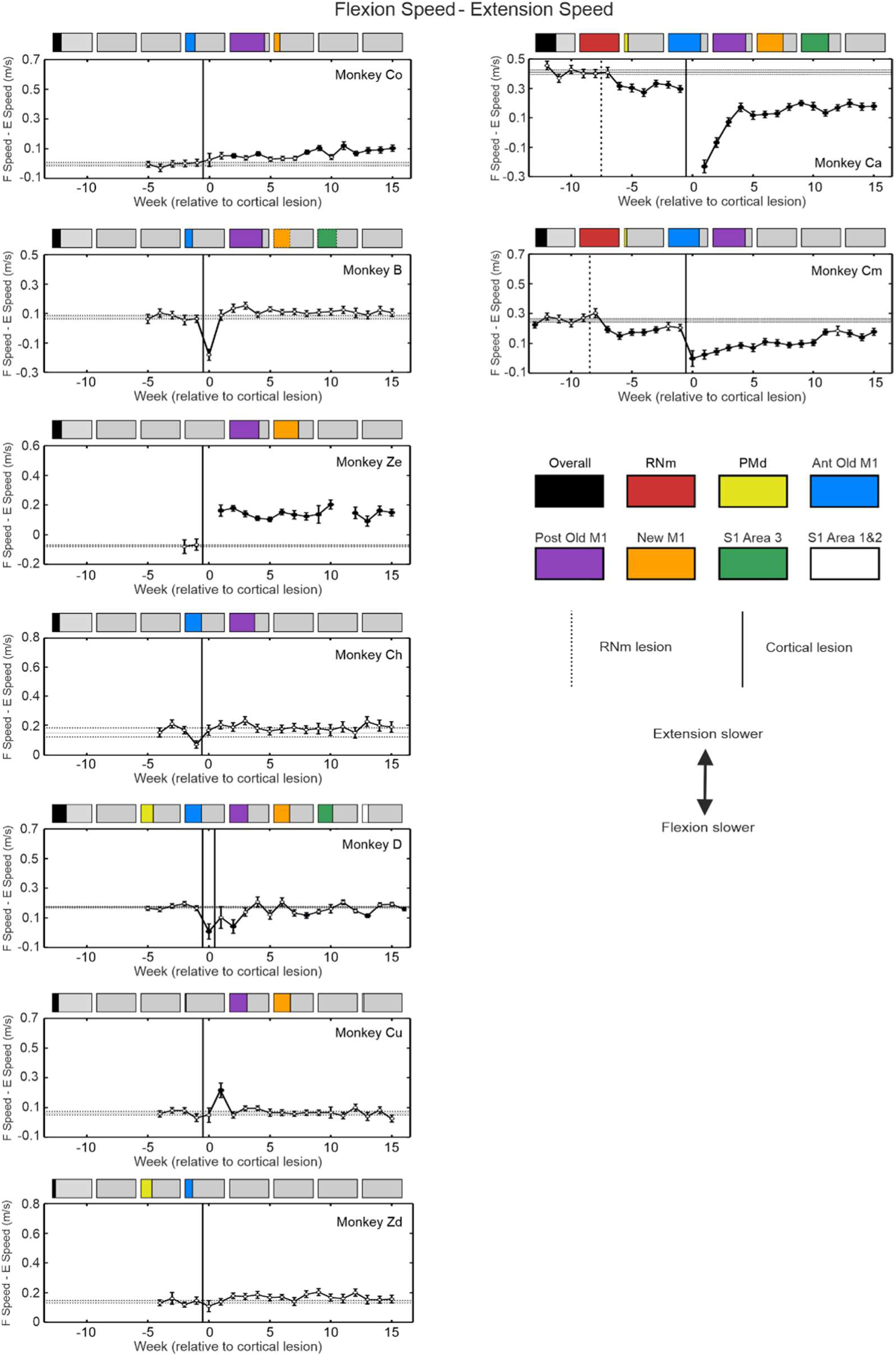
Maximum reaching speed per week compared for elbow flexion vs extension reaching directions. Similar display as Fig. 9, but results shown are the difference between flexion and extension values to appreciate any differences. The y axis range is comparable across all plots between monkeys to allow easier visual comparison. Worse performance in extension is shown by upwards deflection.

Monkeys with lesions almost completely exclusive to M1 generally showed poorer extension reaching performance. This was most evident in Monkey Ze where the speed deficit of elbow extension reaches was persistently greater than for flexion. Greater deficits in AMD^2^ for elbow extension were also seen in all animals with cortical lesions primarily affecting M1 regions, although the magnitude and persistence of this difference varied. No bias was observed in results from Monkey Ch.

A small Anterior Old M1 and PMd lesion caused minimal overall effects to either kinematic measure (Figs. 5 and 6 – Monkey Zd), and there were accordingly small or non-existent elbow flexion-extension differences observed (Figs. 7 and 8).

In Monkey D, the initial lesion of New M1 and Brodmann’s Area 3 produced greater slowing and higher trajectory variability for reaches requiring elbow flexion compared to extension. The second lesion which encompassed PMd, Old M1 and Brodmann’s Area 1 and 2 increased this elbow flexion-extension difference in AMD^2^ initially after the lesion, which recovered slightly more chronically.

### Impact of RNm and Cortical Lesions to Flexion vs Extension Reaching Performance

After RNm lesions alone (dotted vertical line in Figs. 7 and 8), both monkeys showed significantly greater slowing of reaches requiring elbow flexion. There was generally no trend towards worse AMD^2^ in a particular direction in either animal following this lesion (Fig. 7). However, the greater reduction in speed with elbow flexion became exacerbated in both monkeys after cortical lesions (Fig. 8). This is a surprising result, as it is the opposite of the slower reaches requiring elbow extension seen after cortical lesion alone, and which is commonly seen in human stroke survivors. Elbow flexion-extension differences in AMD^2^ after cortical damage were inconsistent, with a brief period of worse elbow extension reaching for Monkey Cm with the smaller cortical lesion, but a larger and more slowly recovering difference towards worse elbow flexion reaches for Monkey Ca. In both animals, reaching speed showed a consistent and long-lasting difference, with slower reaches requiring elbow flexion than extension.

## Discussion

### Relative Differences in Negative Motor Phenotypes Following Focal Cortical Lesions

Lesions that caused damage to Posterior Old and New M1 produced subtle deficits upon observation of the animals, but these were nevertheless clear in kinematic metrics. Table 2 provides a summary of our findings. Both regions have significant corticospinal outputs (Keizer and Kuypers, 1989; Dum and Strick, 1991; He et al., 1993) and CM cell populations (Rathelot and Strick, 2009; Witham et al., 2016), which provide important inputs to upper limb motoneurons in healthy animals. When deprived of these contributions, animals were weak (slowed movements), clarifying the primary facilitatory roles of these parts of M1. Posterior Old M1 additionally has many cortico-reticulospinal outputs (Keizer and Kuypers, 1989; Fisher et al., 2021), unlike New M1. It has been proposed that most drive to motoneurons during a voluntary movement comes from reticulospinal (not corticospinal) neurons (Tapia et al., 2022). It appears that lesions encompassing a larger portion of Posterior Old M1 resulted in a greater reduction of input to motoneurons. This could explain the persistent speed deficits in animals with the most extensive Posterior Old M1 damage, a pattern not seen in animals with less severe Posterior Old M1 involvement but significant damage to New M1 (e.g. Fig. 6 -Monkey Cu).

**Table 2.** Summary of lesion impacts on measures of reaching performance.

| Region of Interest | Effect of Increased Damage to Kinematic Measures | Directional Bias of Impairment | Overt Flexor Synergy |
| --- | --- | --- | --- |
| New M1 | Poorer trajectory variability | Elbow extension | - |
| Posterior Old M1 | More lasting reduced speed | Elbow extension | - |
| Anterior Old M1 | No notable additive negative impact | - | - |
| RNm | Reduced speed after RNm lesion alone<br>Worsened recovery after cortical lesion | Elbow flexion | - |
| PMd, M1, S1 Brodmann's Areas 1, 2 and 3 | Deficits comparable to M1 lesions alone | Elbow flexion | - |

Lesions that produced damage in New M1 and Posterior Old M1 had a negative impact on reaching quality, clearest in monkeys with the largest damage to New M1. Fast-conducting CM cells are found in New M1 and Area 3a (Rathelot and Strick, 2006, 2009). There are also some CM cells in Old M1, but these are slower and generate weaker effects on motoneurons (Witham et al., 2016; Innocenti et al., 2019). Kinematics data gathered from Monkey D in particular highlights the differential effects of lesions to different cell populations (Figs. 5 and 6). Focal lesions to New M1 and S1 Brodmann’s Area 3 negatively affected trajectory variability (but not speed). Subsequent lesions to PMd, Old M1 and S1 Brodmann’s Areas 1 and 2 one week later led to no further impairment in trajectory variability but poorer reaching speed which persisted into the chronic period. Overall, our results suggest that lesions affecting corticoreticular outputs (Old M1) contribute more significantly to weakness because of a greater reduction of overall drive to motoneurons. By contrast, deeper lesions of tissue surrounding the central sulcus (New M1) damage fast corticomotoneuronal outputs and impair the quality of motor control. Some impairment of movement quality is also produced by large, near-exclusive lesions of Posterior Old M1 (e.g. Fig. 5 – Monkey Co); this may be explained by the presence of slower-conducting CM outputs in Old M1.

We suggest that effects following lesions in Monkey D were predominately due to loss of M1 regions, rather than somatosensory and premotor areas. Firstly, deleterious effects on reaching dexterity (increases in AMD^2^ during acute recovery) were comparable following combined lesions of S1 Area 3 and New M1 compared to New M1 alone (Fig. 5 – Monkey Cu). There are no CM cells in Area 3b; the CM cell population found in Area 3a is suggested to target primarily γ-motoneurons involved in fusimotor control (Rathelot and Strick, 2006, 2009). By contrast, CM cells from New M1 project to α-motoneurons at short latencies (Witham et al., 2016). Our findings imply that trajectory variability may be the proximal analogue of dexterity in the hand, which is known to be most significantly mediated by CM cells (Porter and Lemon, 1993).

Secondly, we observed no further initial reductions in reaching speed after combined lesions to PMd and S1 Brodmann’s Areas 1&2 with Old M1, relative to Old M1 lesions alone. This is likely because these areas are predominantly involved in the modulation of M1 output, rather than serving as a primary source of motor commands to α-motoneurons (Widener and Cheney, 1997; Dum and Strick, 2005). This implies that damage to descending pathways, rather than to cortico-cortical connections, is the main determinant of the initial deficit. However, recovery was more complete in the animal with intact PMd and S1, suggesting that some recovery may be mediated by these areas.

This absence of additive negative effects following lesions that involved progressively larger portions of Anterior Old M1 supports the idea that the Area 4s of Hines (1943) has a different role from other M1 subregions. Previous work supported Hines’ idea that it is primarily suppressive (de Barenne and McCulloch, 1941; Boudrias et al., 2010), but we did not find any evidence of disinhibition after Anterior Old M1 lesions. Moderns maps of effects elicited by M1 stimulation show intermixed suppression and facilitation throughout M1 (Boudrias et al., 2010), although suppression is more common for Anterior Old M1. Loss of suppression from Anterior Old M1 could therefore easily by compensated for from other parts of M1. If, as suggested by McCulloch et al. (1946), suppression from Area 4s is partly exerted via cortico-reticulospinal outputs, there would be even more options for recovery, as the primate reticular formation receives widely converging inputs from M1 and pre-motor regions in both hemispheres (Fisher et al., 2021).

### Poorer Recovery with Combined Lesions to RNm and M1 Regions

Non-human primates are a valuable model for translational stroke research (Lin et al., 2023) due to their neuroanatomical and physiological similarities to the human motor system (Porter, 1987; Bortoff and Strick, 1993; Lemon et al., 2002; Lemon, 2008; Decramer et al., 2018; Mu et al., 2023). However, there are clear functional differences in red nucleus outputs between monkeys and humans. RNm cells make an important contribution to upper limb movements in monkeys, particularly those involving the hand (Gibson et al., 1985; Miller et al., 1993; Mewes and Cheney, 1994; Belhaj-Saïf et al., 1998; Van Kan and McCurdy, 2001; Van Kan and McCurdy, 2002a, b). By contrast, the rubrospinal tract is considered vestigial in humans (Hatschek, 1907; Nathan and Smith, 1955; Massion, 1988; Ten Donkelaar, 1988; Basile et al., 2021). With this anatomical background, we hypothesized that RNm lesions combined with lesions to M1 in monkeys would lead to poorer, more human-like recovery.

Our results show that the rubrospinal tract does contribute to recovery of reaching in monkeys following cortical damage. After RNm lesions alone (Fig. 6 after dotted vertical line; Monkey Cm and Ca), reaching movements were significantly slower, likely due to a reduction of rubro-motoneuronal connections that would usually be responsible for facilitating both proximal and distal upper limb muscles (Miller et al., 1993; Belhaj-Saïf et al., 1998). These inputs were irreplaceable by surviving motor areas (ipsilesional M1, supplementary motor area, reticular formation), emphasizing their important contributions to healthy reaching movements in monkey. However, trajectory variability remained unaffected (Fig. 5 after dotted vertical line; Monkey Cm and Ca), possibly because monosynaptic outputs to motoneurons from M1 remained intact. Despite these selective and modest effects of a RNm lesion alone, when combined with cortical lesions the effects were substantial. Combined lesions to RNm, M1 and S1 Area 3 led to severe effects, and the poorest recovery in the entire study (Figs. 5 and 6; Monkey Ca).

We also report a striking better recovery of elbow extension relative to poorer elbow flexion in both animals with RNm lesions (Figs. 7 and 8; Monkey Cm and Ca). This is the opposite to what is typically observed in humans after stroke, where recovery of flexor muscles is favored (Kamper et al., 2003; Baker et al., 2015). It is also unexpected given the known preference for rubrospinal outputs to target extensor muscles more strongly (Mewes and Cheney, 1991; Belhaj-Saïf et al., 1998); we might therefore expect RNm lesions to produce selective deficits in extension. However, after unilateral pyramidotomy rubrospinal neurons exhibit plastic properties and begin to cofacilitate motoneurons innervating flexor muscles and extensor muscles, thereby restoring lost corticospinal inputs to flexors (+340% increased facilitation of flexors, -8% for extensors compared to controls, (Belhaj-Saïf and Cheney, 2000)). This compensatory mechanism may have been compromised following RNm lesions in our animals, with facilitation of extensor muscles but not of flexors, resulting in worse recovery of elbow flexion reaching ability.

Our findings collectively reaffirm that the monkey rubrospinal system is important for restoring input to motoneurons following corticospinal loss, just as has been previously shown in the rat (Ishida et al., 2019). Without functioning rubrospinal outputs, recovery in monkey does become more similar to that in humans, where the rubrospinal tract has been replaced through evolution in favor of an increased reliance on the corticospinal tract.

### No Overt Flexor Synergy Expression Following Focal Cortical Lesions

One suggestion in the literature is that abnormal synergies result from a loss of fine control signals from M1, increasing reliance on motor areas with an innate preference for coding synergistic co-contractions and insufficient fractionated control of shoulder and elbow muscles (e.g. the contralesional cortico-reticulospinal tract, (McPherson et al., 2018)). Lesions to Anterior Old M1 seemed the most appropriate cortical target for generating abnormal flexor synergy based on previous literature. ‘Area 4s’ (found in the anterior region of Old M1) seems capable of exerting powerful suppressive influences over spinal cord motoneuron pools via the cortico-reticulospinal pathway in healthy animals (de Barenne and McCulloch, 1941; Magoun and Rhines, 1946; Engberg et al., 1968; Jankowska et al., 1968; Takakusaki et al., 2001; Boudrias et al., 2010; Du Beau et al., 2012). Ablation of Area 4s was previously described to produce exaggerated stretch reflexes, especially in elbow flexors (Hines, 1936; Hines, 1937, 1943).

We initially hypothesized that isolated damage to Anterior Old M1 would cause impaired suppression of subcortical circuits, resulting in an abnormal flexor synergy pattern. However, our findings did not support this. Eight of the nine animals in this study had lesions that partially involved Anterior Old M1, although the extent of damage varied widely (2-82%). Notably, Monkey Zd regained elbow extension within days after surgery and showed only minimal disruptions in reaching kinematics, despite sustaining damage to approximately 19% of Anterior Old M1 layer V neurons.

One possible explanation is that because our lesions primarily targeted the shoulder, elbow, wrist, and partial hand representations with varying efficacy, sufficient surrounding cortical tissue and descending pathways were spared to permit compensation and prevent the emergence of overt synergy patterns. A similar rationale may apply to the other lesion conditions tested in this study, none of which produced clear synergy patterns. Importantly, all animals retained intact outputs from the supplementary motor area (SMA). Some SMA neurons form monosynaptic connections with spinal motoneurons (Jürgens, 1984) and they exhibit significant plasticity following injury, thereby contributing to functional recovery (Liu and Rouiller, 1999; McNeal et al., 2010).

Early studies which reported positive signs from Area 4s lesions employed lesion techniques where the targeted grey matter was surgically removed. This may have inadvertently damaged underlying white matter, thereby disrupting corticospinal and subcortical control beyond the intended Area 4s. Such more extensive damage could compromise cortical inhibition of subcortical systems and remove options for compensation, and lead to the emergence of abnormal synergies. However, in this study lesions also affected the white matter in some cases (Supplementary Fig. 3A as an example), so that this seems a less likely explanation for lack of synergies in our animals.

To summarize, it seems that the lack of overt flexor synergy expression in our animals was due to focal lesions generating insufficient damage to corticospinal fibers. With sufficient surviving ipsilesional cortex, compensatory strategies involving the contralesional cortex (Cao et al., 1998; Ward et al., 2003) or reticular formation (McPherson et al., 2018; Choudhury et al., 2025) and its indirect connections via spinal cord interneurons (Glover et al., 2026) were probably unnecessary, and the unwanted side effects from using these pathways were avoided.

## Conclusion

This study reveals clear differences between the effects of lesions to various regions of the cortex with different severities. Lesions that damaged greater portions of New M1 had the most significant negative effects on trajectory variability (especially when combined with an Area 3/RNm lesion), as might be expected given its fast CM connections. By contrast, lesions that rendered the most damage to Posterior Old M1 produced longer-lasting deficits in speed; this is also expected, given the extensive cortico-reticular output from this area. Effects of lesions that destroyed large portions of Anterior Old M1 were slight, consistent with the idea that this area plays a distinctively different role in motor control from the other parts of M1, possibly involving suppression. Damage to the RNm alone produced small but persistent effects limited to movement speed, but also markedly accentuated the deficits following subsequent cortical lesions. These results provide new insights into the generation of the multi-faceted phenotype of post-stroke hemiparesis. They reveal that the effects depend on an interplay between descending anatomy of the region damaged, and the ability of other areas to compensate. Strikingly, the effects of a widespread lesion which encompassed approximately a third of the upper limb representation in M1, S1 and pre-motor cortex were no worse than those after more focal lesions to M1. This suggests that many of the consequences of large infarcts often seen from MCA strokes in humans can be understood in terms of component damage to descending pathways, rather than damage to cortico-cortical interactions.

## Acknowledgements

The authors wish to thank Andrew Atkinson, Terri Jackson and Stevie O’Keefe for animal training; Norman Charlton for engineering support; Kathy Murphy, Rocio Palacios O’Connor, Fiona Douglas and Ines Sanchez Garcia for veterinary assistance; and Michelle Waddle for surgical theatre nursing.

