## Supplemental figures for "Arm Control and its Recovery after Selective Lesions of Sensorimotor Cortex and the Red Nucleus: A Kinematic Study in Non-Human Primates"

### Supplementary Material

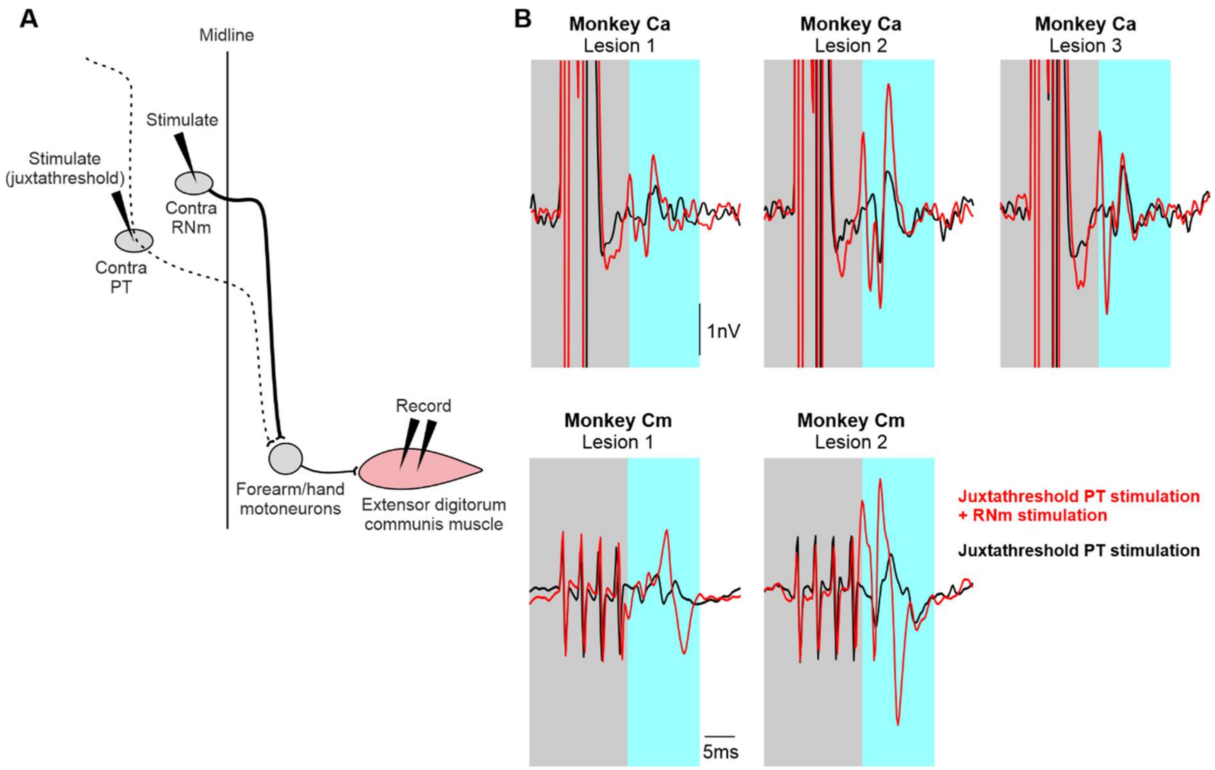


**Supplementary Figure 1. Electrophysiological recording used to guide targeted lesioning of the RNm.** A, schematic of method used to produce RNm lesions in two monkeys. B, electrophysiological recordings from the right extensor digitorum communis muscle for Monkey Ca (top row) and Monkey Cm (bottom row) during RNm mapping. Each column presents EMG traces recorded at the locations where lesions were subsequently made. Motor evoked potentials (blue region) elicited following juxtathreshold left pyramidal tract (PT) stimulation (black traces) or juxtathreshold pyramidal tract stimulation followed by stimulation through the mapping electrode (red traces). Larger responses in red compared to black support that the electrode activated a descending pathway (rubrospinal tract). Grey regions indicate stimulus artefacts.

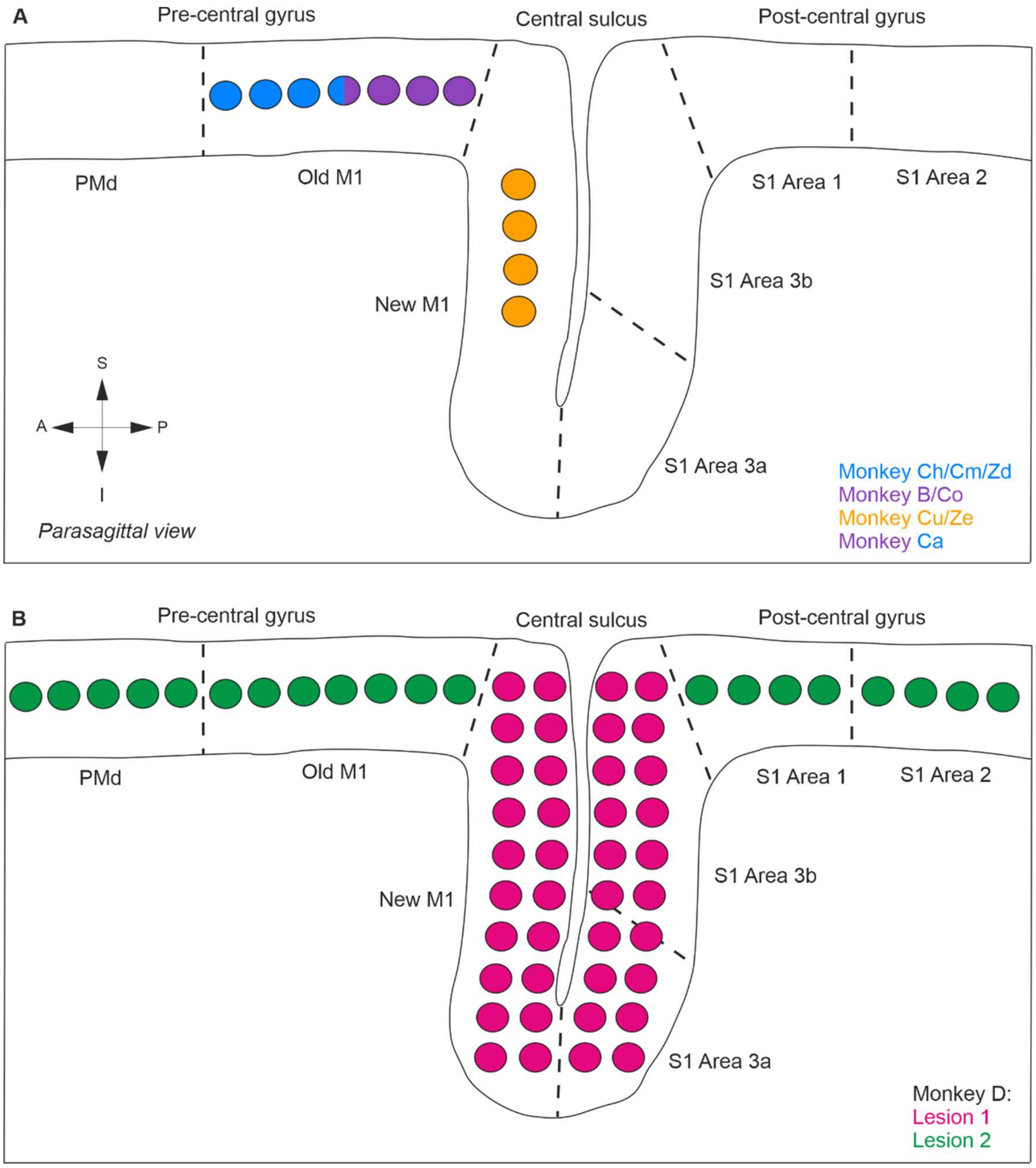


**Supplementary Figure 2. Cortical lesion targets.** A, diagram showing anteroposterior extent of focal lesions performed in different regions of M1. Each circle represents a needle injection site at a single laterality, which extended in a grid 8-19 mm lateral from midline for all animals. Anterior Old M1 lesion (blue); Posterior Old M1 lesion (purple); Entire Old M1 (blue and purple); New M1 lesion (orange). B, diagram of lesion sites performed in Monkey D during two surgeries. Deeper lesions of anterior and posterior banks of the central sulcus (pink); superficial lesions of PMd, Old M1, S1 Brodmann's Areas 1 and 2 (green).

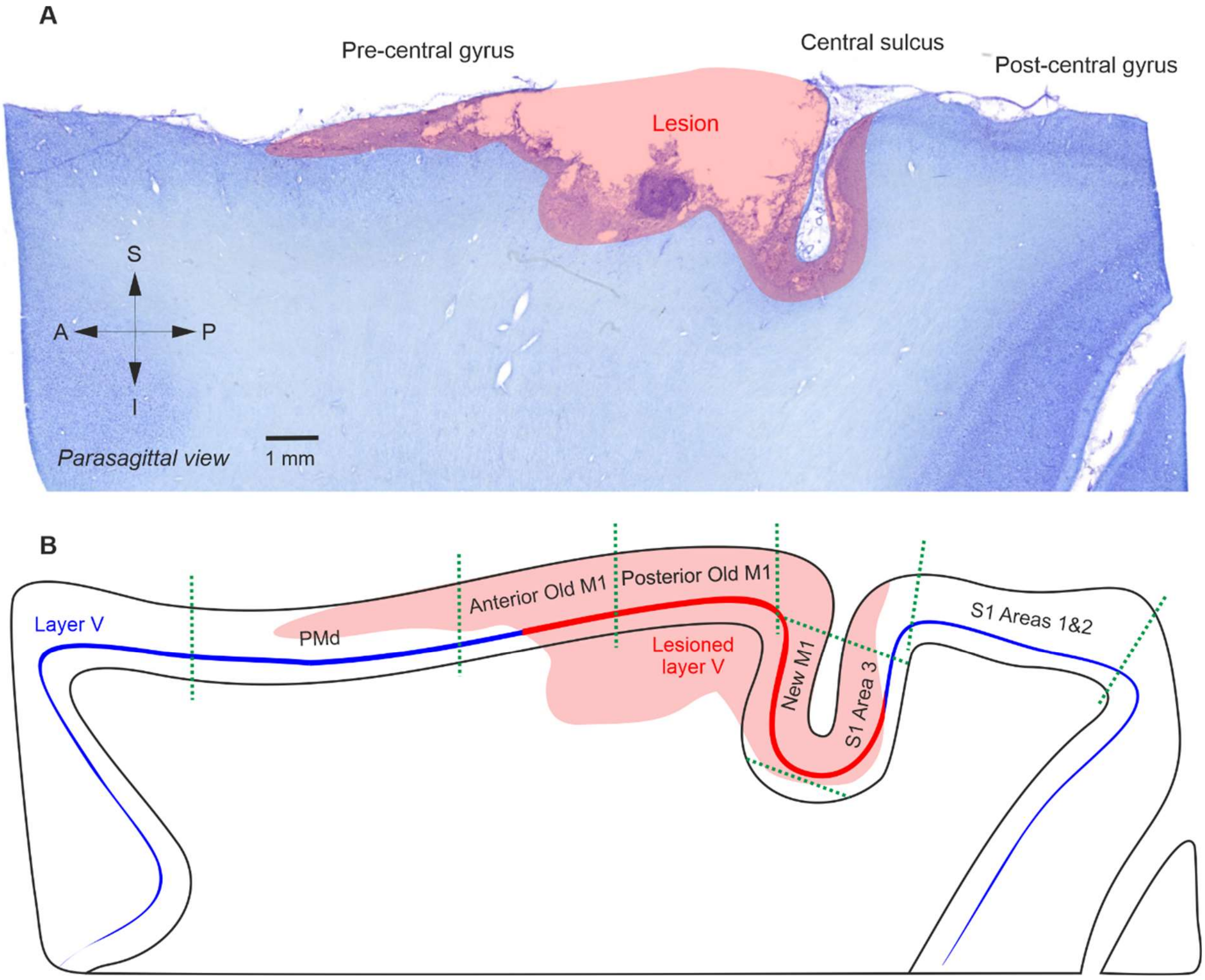


**Supplementary Figure 3. Defining cortical lesion extent with histological staining and physical distance boundaries.** A, example bright field image of a parasagittal section approximately 8 mm from the midline, stained with cresyl violet. The red overlay represents the lesion area. B, corresponding schematic with cortical regions defined using physical distance limits. The red line represents layer V cell layer length that was affected by this lesion; the blue represents unaffected layer V length. For each cortical region, the percentage of layer V cells lesioned were calculated from these two values. *S1 Areas 1&2 = 1-8 mm posterior of the central sulcus (CS) posterior bank; S1 Area 3 superior extent = 1 mm inferior from layer V in S1 Areas 1&2; S1/New M1 inferior extent = most inferior layer V cells in bank of CS; New M1 superior extent = 1mm inferior from layer V in Posterior Old M1; Posterior Old M1 = 1-4 mm anterior to CS; Anterior Old M1 = 4-7 mm anterior to CS; PMd = 7-12 mm anterior to CS.*
